# OsWRKY53 dictates wound responses in rice through fine-tuning cross-talk between PEP and PSK mediated signalling

**DOI:** 10.1101/2025.01.16.633150

**Authors:** Chitthavalli Y. Harshith, Riju Dey, Dipasmit Palchaudhuri, Padubidri V. Shivaprasad

## Abstract

Wounding and tissue damage are major events among the multitude of stresses that plants face in their natural environment. Wound response is a very dynamic event that involves the integration of various regulatory networks culminating in successful wound-induced downstream signalling. Plants depend on endogenously derived molecular signals to initiate wound responses. Transcriptional response is paramount in dictating successful transition of early defense to late growth responses. Here we show the involvement of a WRKY transcription factor (TF) named OsWRKY53, that is responsive to both wounding as well as wound-derived plant elicitor peptide (PEP), OsPep2 treatments. OsWRKY53 is involved in positive regulation of gene expression of OsPep2 responsive genes. OsWRKY53 displays altered DNA occupancy in response to OsPep2 treatment over time correlating with the altered gene expression. Further, OsWRKY53 is involved in simultaneous activation and suppression of OsPep2-responsive and phytosulfokine (PSK) responsive genes, respectively. In agreement with these, mis-expression of OsWRKY53 led to compromised wound responses. Collectively, we establish that OsWRKY53 acts at the intersection of PEP and PSK mediated transition of wound responses and positively regulates the early defense-related gene expression.

## Introduction

Plants encounter a myriad of stresses that involve perturbation of cell wall integrity (CWI). Wounding leads to cell wall integrity breach that triggers a series of molecular signalling events ultimately leading to changes in the gene expression (Harshith et al. 2024; Vega-Muñoz et al. 2020). Wound responses are a culmination of interplay between initial defense outburst and ultimate growth/regeneration responses (Hernández-Coronado et al. 2022; Zhang et al. 2019).

Plants predominantly rely on endogenously derived immunomodulatory cues for signalling cell wall integrity breach (De Lorenzo et al. 2018; Tanaka & Heil 2021; Harshith et al. 2024; Zhai et al. 2024). Damage triggers the activation of genes coding for immunomodulatory peptides that are subsequently processed into bioactive peptides and are broadly referred to as damage associated molecular patterns (DAMPs). Further, DAMPs are perceived by their cognate receptor proteins leading to activation of pattern triggered immune (PTI) responses through MPK signalling cascade. It has recently been shown that in rice, the wound response transition from defense to growth is mediated by two distinct categories of small peptides-PEPs and PSK (Harshith et al. 2024).

PEPs have been reported to be one of the categories of DAMPs that can initiate rapid defense responses immediately upon wounding (Engelsdorf et al. 2018; Harshith et al. 2024). PEPs are 23 aa long bioactive immunomodulatory peptides derived from their non-functional precursors known as PROPEPs (Lori et al. 2015; Yamaguchi & Huffaker 2011). Though PEPs are endogenously derived, the early responses they elicit are similar to responses elicited by patterns of different biological origins in *Arabidopsis* (Bjornson et al. 2021). It has been reported that PEPs can trigger the gene expression very similar to wounding suggesting the general stress activation property of PEPs (Harshith et al. 2024). PEPs are unique in comparison to other stress induced molecular patterns because they are indispensable for wound-induced regeneration in some plants such as tomato (Yang et al. 2024). This indicates the remarkable interplay between PEP-induced defense and PEP-induced growth/regeneration.

Transcriptional changes are the ultimate responses of a signalling event. Wound responses are very dynamic events and involve transcriptional signal relay between initial defense outburst and late growth/regeneration response (Ikeuchi et al. 2017). Such dynamicity is a result of fine-tuned gene expression regulation dictated by transcription factors. The temporal transcriptional response has been shown to be regulated by activation of distinct cis-regulatory elements (CREs) over time post-wounding (Moore et al. 2022). The TF families that regulate late downstream responses leading to wound-induced regeneration or growth have been well studied. These TFs majorly include AP2-domain containing TFs-PLTs, ERFs, WINDs and DOFs (Durgaprasad et al. 2019; Radhakrishnan et al. 2020; Ye et al. 2020; Iwase et al. 2013; Zhang et al. 2022). Also, for the earliest transcriptional changes post-wounding, MYC family of TFs that are downstream to JA signalling have been attributed (Walley et al. 2007). However, the TFs that potentially act at the intermediate stage right during initial outburst and during the onset of signal transitioning over time are not investigated. Among the predominantly enriched CREs post-wounding, W-box elements recognized by WRKY group of TFs are present in the promoters of the genes that rapidly respond to wounding (Moore et al. 2022; Rushton et al. 2002). The absence of wound-induced regeneration in rice suggests possible involvement of distinct TFs that can potentially dictate signal transition by changing the gene regulatory network leading to quick transition to growth phase post-wounding.

WRKY family of TFs are plant-specific and are predominantly involved in regulation of various stress responses. WRKYs are involved in very early signalling upon PTI activation as suggested by predominant enrichment of their CREs in the genes that are rapidly induced upon perception of various immune triggers (Bjornson et al. 2021). Further, it has been shown that WRKYs usually act in a sub-regulatory network leading to both auto as well as cross regulation of other WRKYs (Birkenbihl et al. 2018). This regulatory network translates to determination of nature of temporal gene expression upon perception of various stresses. WRKY proteins possess conserved DNA binding domains with the conserved WRKYGQK sequence (Yamasaki et al. 2005). WRKYs are classified into three major groups based on the number of DNA binding domains and the kind of zinc-finger motifs (Brand et al. 2013). Main groups are further subdivided into sub groups based on additional specificities and domains involved in interaction with other proteins (Chi et al. 2013). Functions of WRKYs are not restricted to stress response, they are required for the regulation of growth, development, senescence, germination, etc (Wang et al. 2023; Zentgraf et al. 2010). The role of WRKYs in regulation of transition of defense to growth responses is not understood and it is possible that WRKYs are involved in this owing to their versatility.

PEP signalling initiates after perception of PEP by cognate receptor protein, PEPR (Yamaguchi et al. 2006; Shinya et al. 2018). The receptor complex activation leads to initiation of MPK signalling cascade (Harshith et al. 2024) eventually activating the TFs that mediate gene expression changes. Group I WRKYs have been attributed to be direct substrates of MPK protein as they possess D-domain required for interaction with MPKs (Ishihama & Yoshioka 2012). These WRKYs possess SP cluster in the N-terminal region in the vicinity of D-domain (Ishihama & Yoshioka 2012). Serine residues in SP cluster undergo phosphorylation mediated by MPKs leading to activation of WRKYs. Many group I WRKYs have been identified to be substrates of MPKs in rice and have been attributed to plethora of functions including regulation of disease resistance. Among the well-studied group I WRKYs in rice, OsWRKY53 seems to be very versatile as it is involved in regulation of insect herbivore responses, disease resistance, grain size, grain yield, seed germination, brassinosteroid signalling, cold stress, salt tolerance and leaf senescence (Tang et al. 2022; Xie, Ke, et al. 2021; Tian et al. 2021; Hu et al. 2015; Chen et al. 2024; Xie, Li, et al. 2021; Yoo et al. 2014; Chujo et al. 2014; Chujo et al. 2007). OsWRKY53 is involved in regulation of both growth and defense (Hao et al. 2022; Meng et al. 2024). OsWRKY53 has also been shown to be upregulated upon wounding (Hu et al. 2015; Harshith et al. 2024). The global transcriptional regulation dictated by OsWRKY53 has not been understood in the context of wounding or PTI responses.

In this study, we have identified a WRKY TF, OsWRKY53 responsive to wounding, PEP treatment as well as PSK treatments. During homeostasis, OsWRKY53 was expressed at basal level and regulated genes involved in some of the metabolic processes. Treatment of rice plants with OsPep2 directly altered the chromatin occupancy of OsWRKY53 by both intensifying the occupancy and changing the regions occupied over time as observed in chromatin immunoprecipitation followed by sequencing (ChIP-seq) experiments. Further, OsWRKY53 directly contributed to the expression of a proportion of OsPep2 responsive genes by directly occupying their promoters. OsWRKY53 was autoregulatory in nature and contributed to cross-regulation of many other WRKYs. OsWRKY53 also regulated gene expression post-wounding by positively regulating early wound-responsive genes and negatively regulating the late responsive genes. The transcriptome of the wound responses at 12 h post wounding in Oswrky53 knock-out (KO) resembled that of OsPSKR overexpression (OE) suggesting a contrasting role between OsPSKR and OsWRKY53 mediated gene expression. Further, in the Oswrky53 KO plants, PSK-responsive genes were constitutively mis-expressed and they were actively suppressed when OsPep2 signalling was activated. This suppression of PSK responses was dependent on OsWRKY53 indicating that OsWRKY53 contributes to early defense signalling downstream to wounding and OsPep2 treatment by simultaneous suppression of growth signalling dictated by PSK.

## Materials and methods

### Plant materials and growth conditions

*Oryza sativa indica* Pusa Basmati 1 (PB1) variety was used in this study. The rice seeds were germinated in half-strength MS medium in dark before transferring to light. Fourteen-day old seedlings were transferred to greenhouse and grown at 28°C under natural day-night cycle.

### Plasmid construction and cloning

To generate CRISPR knock-out construct, guide RNA fragments – gRNA1 and gRNA2 were inserted into pRGEB32 vector (Xie et al. 2015) using BsaI enzyme sites. To generate OsWRKY53-GFP constructs used for ChIP experiments, OsWRKY53 promoter region, 2.3 kb in length was amplified from PB1 genomic DNA. This promoter fragment was inserted into pCAMBIA1380 (Genbank accession no. AF234301.1) between BamHI and SalI sites. Further, OsWRKY53-CDS and GFP coding regions were inserted between SalI and HindIII sites in the vector containing OsWRKY53 promoter. All the above mentioned vectors were mobilized into Agrobacterium strain – LBA4404 with pSB1 (Sridevi et al. 2003).

### Rice transformation

Rice transformation was performed as described previously (Sridevi et al. 2003; Swetha et al. 2018). Rice seeds of PB1 were dehusked and surface sterilized using washes with 70% ethanol, 4% bleach and 0.1% mercuric chloride. Sterilized seeds were placed on callus induction media and maintained in dark for 21 days. The embryogenic calli obtained were used for transformation with desired constructs using LBA4404 (with pSB1) strain of *A. tumefaciens*. After transformation, calli were selected on media containing hygromycin antibiotic. The selected calli were transferred to regeneration media and the shoots derived were transferred to half-strength MS media for rooting before transferring to soil.

### Leaf wounding experiments

All the wounding experiments mentioned in the text were performed on six-week-old rice leaves. Leaf blades were injured by punching the edges at 2 cm interval using a 3 mm punching equipment without damaging the midrib of the leaves (Venu et al. 2010). Wounded leaves were collected at the corresponding time points along with their unwounded controls and snap frozen before processing for transcriptome profiling.

### Peptide treatment

Six-week-old rice leaves were used for all the peptide treatment experiments. Rice leaves were cut into strips of 2 cm each and were maintained in sterile water overnight to wash away the residual wound signals as described previously (Schwessinger et al. 2015). Leaf strips were treated with OsPep2 or PSK peptides of 1 µM concentration (custom made from Lifetein) and collected at the corresponding time points mentioned in the text. The samples were snap frozen for further experiments.

### RT-qPCR experiments

For RT-qPCR experiments, first strand cDNA was synthesized from the extracted RNA using Thermo RevertAid first strand cDNA synthesis kit following manufacturer’s instructions. The obtained cDNA was used for qPCR experiments. SYBR green master mix (Solis Biodyne-5X HOT Firepol Evagreen qPCR master mix) was used for qPCR reaction. *OsActin* was used as internal reference. The primers used are mentioned in the Table S14.

### Transcriptome analysis

For RNA-sequencing experiments, six-week-old plants were used. For general sequencing and wounding experiments, entire leaf blades were used. For OsPep2 and PSK treatment experiments, leaf strips were used. Samples were snap frozen after collection at the indicated time points. The ground powder was used for RNA extraction using Trizol method. Obtained RNA was poly A enriched for library preparation. Library preparation was carried out using NEBNext® Ultra™ II Directional RNA Library Prep kit (E7765L) according to manufacturer’s protocol. Subsequent sequencing was performed in paired-end (2x100bp) manner on NovaSeq 6000 platform. RNA sequencing analysis was carried out as described previously. For all the samples, an average of 30 million paired-end reads were obtained. Trimmomatic tool was used for adapter trimming (Bolger et al. 2014). After adapter trimming, reads were aligned to the genome (IRGSP1.0) of rice using HISAT2 (Kim et al. 2015). Further, Cufflinks was used to obtain the gene expression level and the results are depicted through plots generated using R (Trapnell et al. 2014).

### GO analysis

GO analysis for various categories of genes was performed using ShinyGO v0.80 platform. The gene IDs used were obtained from RAPDB. The GO chart was obtained for parameters-biological processes with FDR cut-off of p-value – 0.05.

### Southern blotting

Southern blotting assay was performed as mentioned previously (Pachamuthu et al. 2022). Total DNA extraction was performed using CTAB method of DNA extraction. Around 10 µg of DNA was digested with the enzymes indicated. Digested DNA was run and resolved on 0.8% gel in 1XTBE. After gel running, DNA was transferred to Zeta probe nylon membrane (Biorad) using capillary transfer method. Membrane was crosslinked using UV after transfer. The DNA probe used was DNA coding for hygromycin resistance gene. The probe was amplified using PCR from the plasmids containing this DNA. Further, the probe was internally labelled with [α-P32] dCTP (BRIT India) using Rediprime labelling kit (GE healthcare). The labelled probe was used for hybridization of the membrane and the membrane was subjected to phosphor imaging. The membrane was scanned for phosphor scanning using Typhoon scanner (GE healthcare).

### ChIP-sequencing and analysis

OsWRKY53-GFP plants were used for ChIP experiments. Six-week-old leaf strips were subjected to the treatments indicated in the text followed by crosslinking. Further, 1 g of crosslinked tissue was used for each sample and processed as described earlier (Saleh et al. 2008; Hari Sundar G et al. 2023). After chromatin shearing, the samples were incubated overnight with GFP trap beads from chromotek. After washes and elution of DNA, library preparation was performed using NEBNext Ultra II DNA library prep kit (NEB E7103) following manufacturer’s instructions. The sequencing was performed on Novaseq 6000 platform.

Obtained reads were used for analysis. Adapter trimming was performed using Cutadapt and the trimmed reads were mapped to IRGSP1.0 genome using Bowtie 2 (Langmead & Salzberg 2012; Martin 2011). The parameters used for bowtie: -v 1 -k 1 -y -a -best – strata. PCR duplicates were removed and converted into bigwig tracks using subtract option in deepTools using WT sample as control (Ramírez et al. 2014). For generating meta plots, ComputeMatrix (deepTools) was used and the plots were generated using plotprofile. Narrow peaks were called using MACS2 with respect to WT samples (Zhang et al. 2008). Obtained peaks were analysed and the replicates were merged using BEDTools (Quinlan & Hall 2010). The region annotation of the origin of peaks was done through ChIPseekeR (Yu et al. 2015). The overlap Venn diagrams were generated through Intervene (Khan & Mathelier 2017).

### Protein extraction and immunoblotting

The leaf samples derived from six-week-old plants were subjected to OsPep2 or PSK treatment as indicated above. After the collection at time points as indicated in the main text, the samples were ground using liquid nitrogen in mortar and pestles. The proteins were extracted using TCA method as described previously (Nair et al. 2020). Briefly, the proteins were initially precipitated using 10% TCA followed by 80% methanol with 0.1 M ammonium acetate. Further, proteins were extracted from the obtained pellet using the extraction buffer containing: buffered phenol, 0.876 M sucrose, 0.1 M Tris-Cl pH-8.0, 0.75 M β-ME and 2% SDS. The upper phase was then used to pellet the proteins using 100% methanol followed by 80% acetone. The obtained pellet was dissolved in 2X Lamelli sample buffer and loaded on gels. The western blotting using GFP antibody was performed as described previously (Harshith et al. 2024).

### Phos-tag assay

For detection of phosphorylation status of OsWRKY53 upon OsPep2/PSK treatment, the protein samples obtained using the above-mentioned procedure were run on Mn^2+^ - Phos-tag gels. The resolving gels were prepared using 0.5 mM Phos-tag Acrylamide AAL-107 reagent from sigma (TA9H9A175FCA) along with 10 mM MnCl_2_. Before setting up transfer, the gels were washed in the transfer buffer containing 1 mM EDTA. Further, the western blotting was performed as described above.

## Results

### OsWRKY53 is an early responder to wounding, PEP and PSK treatments

Transcriptional responses downstream to wounding overlap with that of responses to wound-derived OsPep2 and OsPep3 treatments (Harshith et al. 2024). In wounded rice leaves, signal transition from initial outburst of defense to later growth responses relied on a nodal receptor protein OsPSKR involved in wound-induced PSK perception (Harshith et al. 2024). We hypothesized that such a response necessitates the requirement of proteins having the ability to participate in signalling downstream to both defense and growth pathways. Since OsPSKR knock-out (KO) plants were severely compromised in overall growth of the plants, we hypothesized that the genes involved in wounding as well as in OsPSKR mediated signalling might be important for temporally regulating the transition of wound responses. In order to identify such candidates, we considered all the genes that were significantly upregulated upon wounding as well as wound-derived PEP treatments at an early time point (30 min) that captures genes involved in defense outburst. Among the 165 candidate genes we obtained, 39 of them overlapped with 381 significantly downregulated genes in OsPSKR KO (Fig. 1A). These 39 genes exhibited positive response to both wound and PSK mediated pathways. These included various defense response-associated genes such as WRKYs, MPKs, JAZ repressors, bHLH TFs, those involved in JA signalling and in transcriptional regulation (Fig. S1A and S1B, Table S2). Interestingly, most of these 39 genes were also upregulated in OsPSKR OE indicating the specificity and importance of these genes downstream to OsPSKR mediated signalling (Fig. S1C, Table S2). To explore if they also respond to PSK treatment, we performed PSK treatment in WT (PB1 variety of *Oryza sativa indica*) leaves. Surprisingly, a majority of these 39 genes showed a very early response to PSK treatment (Fig. S1C), suggesting that they are indeed downstream to both PEP induced defense activation and PSK induced growth activation.

**Figure 1:**
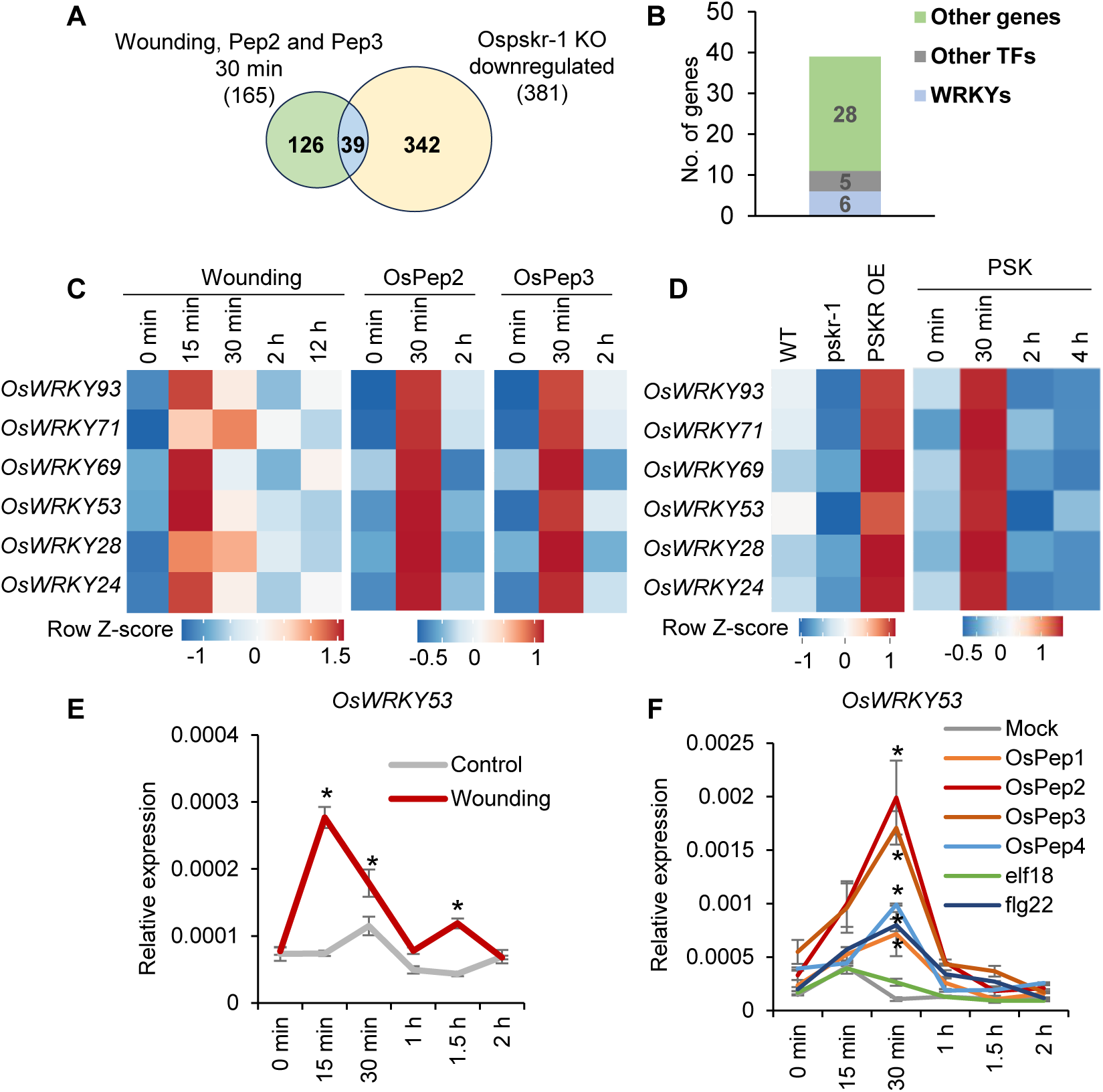
OsWRKY53 is an early responder to wounding, PEP and PSK treatments. **(A)** Overlap of genes between wounding, OsPep2 and OsPep3 treatments upregulated at 30 min with those that are downregulated in case of Ospskr-1 KO, Log_2_FC ≥ 1.5 for upregulated and Log_2_FC ≤ -1.5 for downregulated, p<0.05. **(B)** Number and type of genes downstream to OsPSKR and responsive to wound and wound-derived peptide treatments. **(C)** Heatmap showing the expression of WRKY TFs upon wounding and wound-derived PEP treatment. **(D)** Heatmap showing the expression of WRKY TFs in OsPSKR mis-expression lines and PSK treatment. For **(C** and **D)**, gene expression is represented as row Z-score of FPKM obtained from average of replicates in RNA-seq experiments. **(E)** RT-qPCR analysis of expression of Os*WRKY53* upon wounding. **(F)** RT-qPCR analysis of expression of Os*WRKY53* upon different peptide treatments. *OsActin* served as internal controls for RT-qPCR. n - 3 for all qPCR experiments, error bars indicate standard error of the mean (SEM), pairwise Student’s *t*-test with respect to corresponding untreated samples; ∗p < 0.05.

Since TFs play important regulatory roles in determining the outputs of signalling events, we narrowed down to 11 TF coding genes. Six of them belonged to WRKY family of TFs (Fig. 1B). The other TFs included 3 bHLH TFs, a MYB TF and a NAC TF (Fig. 1B and S1B). Since WRKY TFs are predominantly involved in stress responses with a variety of roles (Javed & Gao 2023), we considered them for further investigation. All six WRKYs showed a very rapid induction both upon wounding as well as with wound-derived OsPep2 and OsPep3 treatments (Fig. 1C, Table S1). Most importantly, all these WRKYs were upregulated in OsPSKR OE lines and also showed a very rapid response to PSK treatment indicating their involvement in both PEP and PSK signalling pathways (Fig. 1D, Table S1). Among the six WRKYs that were showing response, OsWRKY53 and OsWRKY69 showed pronounced increase in comparison to others. OsWRKY53 was previously implicated in insect herbivore responses, disease resistance, brassinosteroid signalling pathway, negative regulation of abiotic stress tolerance, etc. (Tang et al. 2022; Xie, Ke, et al. 2021; Yu et al. 2023; Tian et al. 2021; Hu et al. 2015). OsWRKY53 belongs to group-I of WRKYs, members of which are predominant substrates of MPK signalling downstream to PTI activation (Ishihama & Yoshioka 2012). These attributes of OsWRKY53 led us to explore its role in the context of wound responses mediated by small peptides.

OsWRKY53 showed a very rapid response to wounding transcriptionally with increased expression at 15 min time post-wounding (Fig. 1E). Since OsWRKY53 showed upregulation upon OsPep2, OsPep3 treatment as well as with PSK, we checked if this response is specific. In a RT-qPCR experiment involving plant-derived PEPs (OsPep1-4), bacterial flg22 and elf18 peptides, upregulation of OsWRKY53 was very high and rapid in case of OsPep2 and OsPep3 treatments, but only marginally with OsPep1, OsPep4 and flg22 treatments (Fig. 1F). OsWRKY53 was not responsive to elf18 as expected, as rice lacks receptor for elf18 peptide (Schwessinger et al. 2015) indicating specificity of OsWRKY53 in PTI responses (Fig. 1F). These results indicated that OsWRKY53 responds to various peptides for which there are receptors to activate PTI and the response is stronger upon wound derived OsPep2 and OsPep3 treatments.

### OsWRKY53 contributes to gene expression control during homeostasis

To understand the specific roles of OsWRKY53 in wound-derived peptide-mediated signalling, we generated two guide RNAs (gRNA) to specifically edit OsWRKY53 coding region. The two gRNAs-gRNA1 and gRNA2 were designed to target exon-2 and exon-3, respectively (Fig. 2A). We obtained two independent lines with a single nucleotide insertion leading to premature stop codon in the gRNA1 targeted line which will be referred to as KO1 henceforth (Fig. 2B-D). The line targeted by gRNA2 had a deletion of 5 nucleotides resulting in premature stop codon which will be referred to as KO2 (Fig. 2B-D). The KO plants did not exhibit drastic phenotypic changes in comparison to WT non-transgenic plants (Fig. 2E). In order to check the role of OsWRKY53 in contribution to gene expression status during homeostatic conditions, we performed transcriptome profiling using both the KO lines along with WT control. We observed a total of 3495 differentially expressed genes (DEGs) in KO1 and 2118 DEGs in KO2 after imposing a stringent cut off of Log_2_FC ≥ 1.5 for upregulated genes and Log_2_FC ≤ -1.5 for downregulated genes (Fig. 2F, Table S3). More than 75% of the DEGs were common between both the lines suggesting that both biologically distinct lines were affected in similar pathways (Fig. 2F, Table S3). The magnitude of KO1 was severe in perturbation of gene expression in comparison to KO2, and this might be due to retention of N-terminal stretch in the truncated protein. The gene ontology (GO) analysis of the DEGs in both the lines indicated that the upregulated genes were predominantly involved in various metabolic processes and the downregulated genes were involved in rRNA metabolism and biosynthetic processes (Fig. 2G and H). In agreement with a previous report indicating negative regulation of sugar transporter gene OsSWEET1b by OsWRKY53 (Chen et al. 2024), we observed upregulation of this gene in both the KO lines (Fig. 2I). Also, OsWRKY53 has been shown to be involved in positive regulation of brassinosteroid (BR) signalling (Tian et al. 2017). We noticed that OsBRD2, involved in BR biosynthesis, was downregulated in both the KO lines (Fig. 2J). Together, these results suggest that the basal expression of OsWRKY53 is important for the regulation of gene expression during homeostasis.

**Figure 2:**
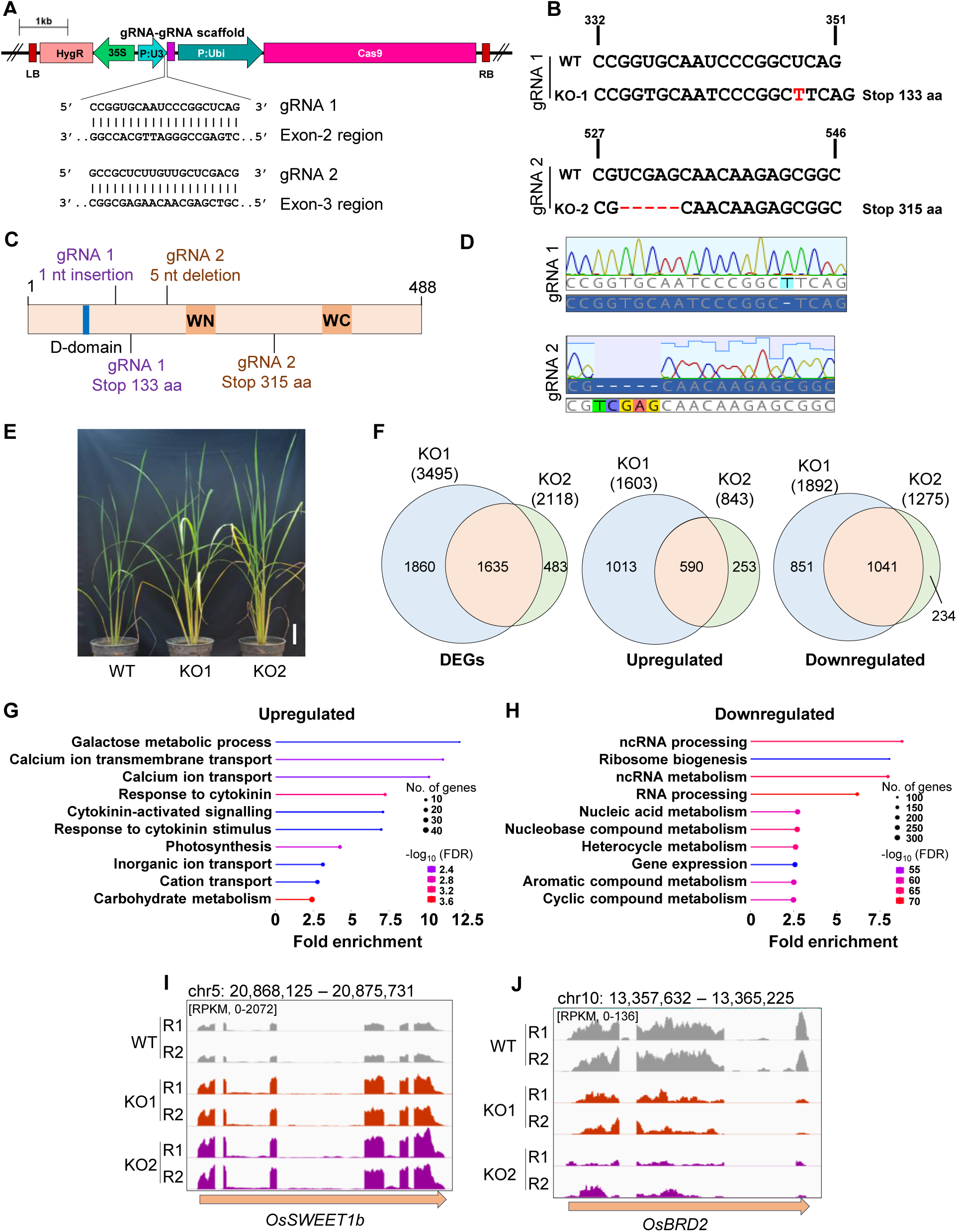
OsWRKY53 contributes to gene expression control during homeostasis. **(A)** T-DNA map depicting the guide RNA constructs used for generating OsWRKY53 KO plants. **(B)** Depiction of mutants obtained after CRISPR editing. **(C)** Schematic depicting domains of OsWRKY53 and showing location of premature stop codons in two KO lines. **(D)** Chromatograms showing sequences of OsWRKY53 coding regions in edited regions of KO lines. **(E)** Phenotypes of OsWRKY53 KO lines (12-week-old). Scale-8 cm. **(F)** Overlap of DEGs between two KO lines. UP - Log_2_FC ≥ 1.5, p<0.05, DOWN - Log_2_FC ≤ -1.5, p<0.05 DEGs - Differentially expressed genes. **(G)** GO analysis of genes significantly upregulated in both the KO lines of OsWRKY53. **(H)** GO analysis of genes significantly downregulated in both the KO lines of OsWRKY53. **(I-J)** IGV screenshots depicting the expression pattern of *OsSWEET1b* and *OsBRD2* in KO lines compared to WT.

### OsWRKY53 occupies genic and distal regions of a subset of genes and contributes to their expression

In order to understand the downstream target genes regulated by OsWRKY53, we generated GFP tagged OsWRKY53 lines where the transgene was driven by native promoter of OsWRKY53 (Fig. S2A). We obtained two independent transgenic lines with single copy T-DNA insertion events (Fig. S2B). We used line #7 for subsequent experiments and analysis (Fig. S2B). The plants displayed phenotypes similar to increased brassinosteroid responsive plant phenotypes (Fig. S2C) reported earlier for overexpression of OsWRKY53 (Tian et al. 2017). After confirming transgenic protein expression through western blot analysis (Fig. S2D), we performed ChIP-seq to identify the downstream target genes during homeostasis. We obtained 3912 peaks where OsWRKY53 displayed clear binding (Fig. 3A). The peaks predominantly originated from distal intergenic regions (Fig. 3B). A little over 20% of the peaks obtained were from proximal and distal promoter regions of protein coding genes (Fig. 3B). In order to ascertain the magnitude of peaks originating from the gene and gene regulatory regions, we intersected the obtained peaks with that of gene body regions (+2 kb promoter and terminator regions). Among the 3912 peaks obtained, 732 peaks originated from the genic regions (Fig. 3C). These peaks were distributed over promoters, terminators and gene body regions (Fig. 3D). Out of these, a total of 315 peaks originated from promoter regions, while gene bodies intersected with 201 peaks, and 286 were mapped to terminator regions (Fig. 3E). Binding of OsWRKY53 to these distinct regions showed positive regulation of expression of some of the genes (Fig. 3F-H). For instance, OsWRKY53 occupied the promoter of the gene *Os09g0491596* and the expression was downregulated in both the KO lines (Fig. 3F). Similarly, OsWRKY53 occupied the gene body and terminator of gene *Os03g0349000* and OsGRX2, respectively, mediating the positive regulation of expression of these two genes (Fig. 3G and H). We then compared the expression of genes directly occupied by OsWRKY53 and found that a few genes were significantly downregulated in KO lines, indicating that they might be under positive regulation of OsWRKY53 (Fig. S2E and G). On the other hand, a few genes were upregulated in KO lines, indicating these genes might be negatively regulated by OsWRKY53 (Fig. S2F and S2H, Table S4). This suggests that OsWRKY53 has specific targets during homeostasis, likely involved in development and growth.

**Figure 3:**
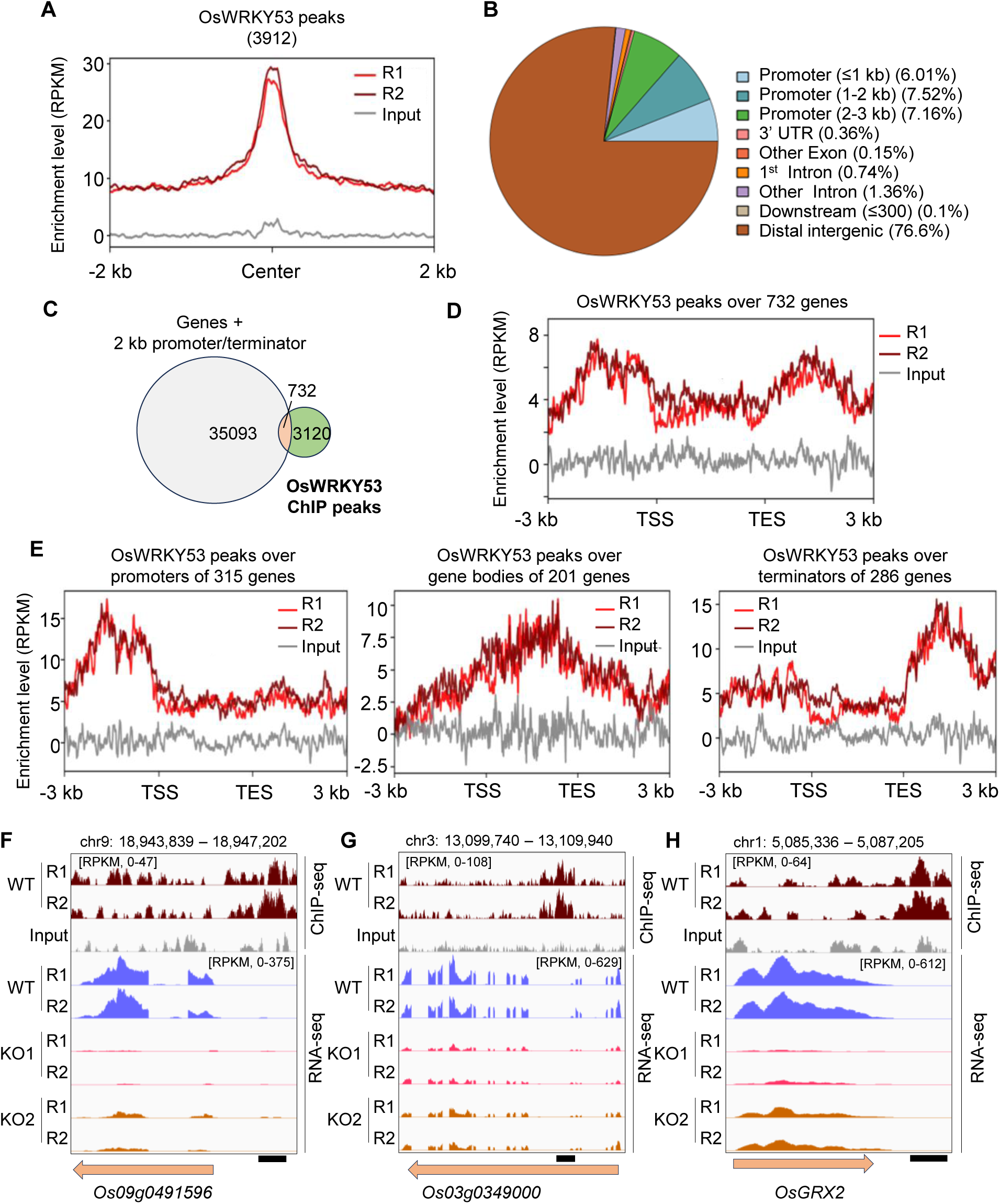
OsWRKY53 occupies genic and distal regions of a subset of genes and contributes to their expression. **(A)** Metaplot showing the enrichment of OsWRKY53 during homeostasis. **(B)** Pie chart depicting distribution of OsWRKY53 bound regions. **(C)** Overlap between OsWRKY53 bound peaks and genic regions. For each gene, 2 kb promoter and 2 kb terminator were considered. **(D)** Metaplot showing distribution of OsWRKY53 peaks over 732 genes that overlapped with OsWRKY53 peaks. **(E)** Specificity of occupancy of OsWRKY53 over promoters, gene-bodies and terminators of genes overlapping with OsWRKY53 peaks. **(F-H)** IGV screenshots showing the expression of genes influenced by direct binding of OsWRKY53. R1 and R2 are replicates. Input represents the DNA before antibody-based enrichment. TSS – transcription start site, TES – transcription end site, Centre-Centre of each peak. The arrows below IGV images represent genes and the black bands represent OsWRKY53 binding regions.

### OsWRKY53 responds to OsPep2 treatment and shows altered occupancy over chromatin

To explore the specific signalling role of OsWRKY53 in the modulation of gene expression downstream to PEP perception, we treated six-week-old rice leaves with OsPep2 and performed ChIP-seq at two different time points to understand signalling both at early and late time points. The overall number of peaks obtained post OsPep2 treatment changed marginally over time. The mock treated samples showed a total of 3742 OsWRKY53 enriched peaks, whereas 30 min post treatment the number of peaks increased to 4387 and reverted back to 3984 at 2 h post treatment (Fig. S3A). We further probed if the distribution of peaks across genomic regions were altered in response to OsPep2 treatment, and observed that occupancy was marginally increased on promoter regions at 30 min time point (Fig. 4A). The peaks originating from genic regions increased at 30 min with around 400 additional peaks in comparison to mock treatment and 2 h time points (Fig. S3B-D). These peaks were distributed over promoters, terminators and gene bodies (Fig. 4B-D). The intensity of peaks occupying promoters increased substantially at 30 min time point when compared to mock, suggesting a likely contribution of OsWRKY53 in the promotion of PEP-responsive gene expression (Fig. 4C). To tease apart the specific and generic roles of OsWRKY53, we intersected the peaks obtained at mock as well as across time points upon OsPep2 treatment and observed that 1728 peaks were persistent across the treatment regime. These peaks showed sharp increase in binding at 2 h time point (Fig. 4E-F). Among the unique peaks, 897 were obtained in mock treated conditions, increased to 1502 and 1576, respectively, at 30 min and 2 h post treatments (Fig. 4E-F). The enrichment level of mock unique peaks was just marginally different from the rest (Fig. 4F and S3E). However, the intensity of 30 min unique peaks (1502 peaks) increased in response to OsPep2 treatment (Fig. 4F and S3E). The response of unique peaks at 2 h point (1576 peaks) was completely distinct from the rest, having bimodal peaks, suggesting altered chromatin occupancy (Fig. 4F and S3E). We observed that among the persistent peaks, there were around 374 peaks which were longer than 500 nt (broad peaks) and showed similar enrichment profile across treatment regime (Fig. 4G and S4A). We also noticed that, at 2 h post treatment, a total of 167 peaks were longer than 500 nt and these peaks were completely unique to 2 h OsPep2 treatment (Fig. 4H and S4A). Such an enrichment of broad peaks was observed even 2 kb away from the centre of the peaks, unlike observed across plant TFs (Fig. 4F). All these indicate a dynamic binding property of OsWRKY53 in response to wounding and OsPep2 treatments, likely to mediate gene expression.

**Figure 4:**
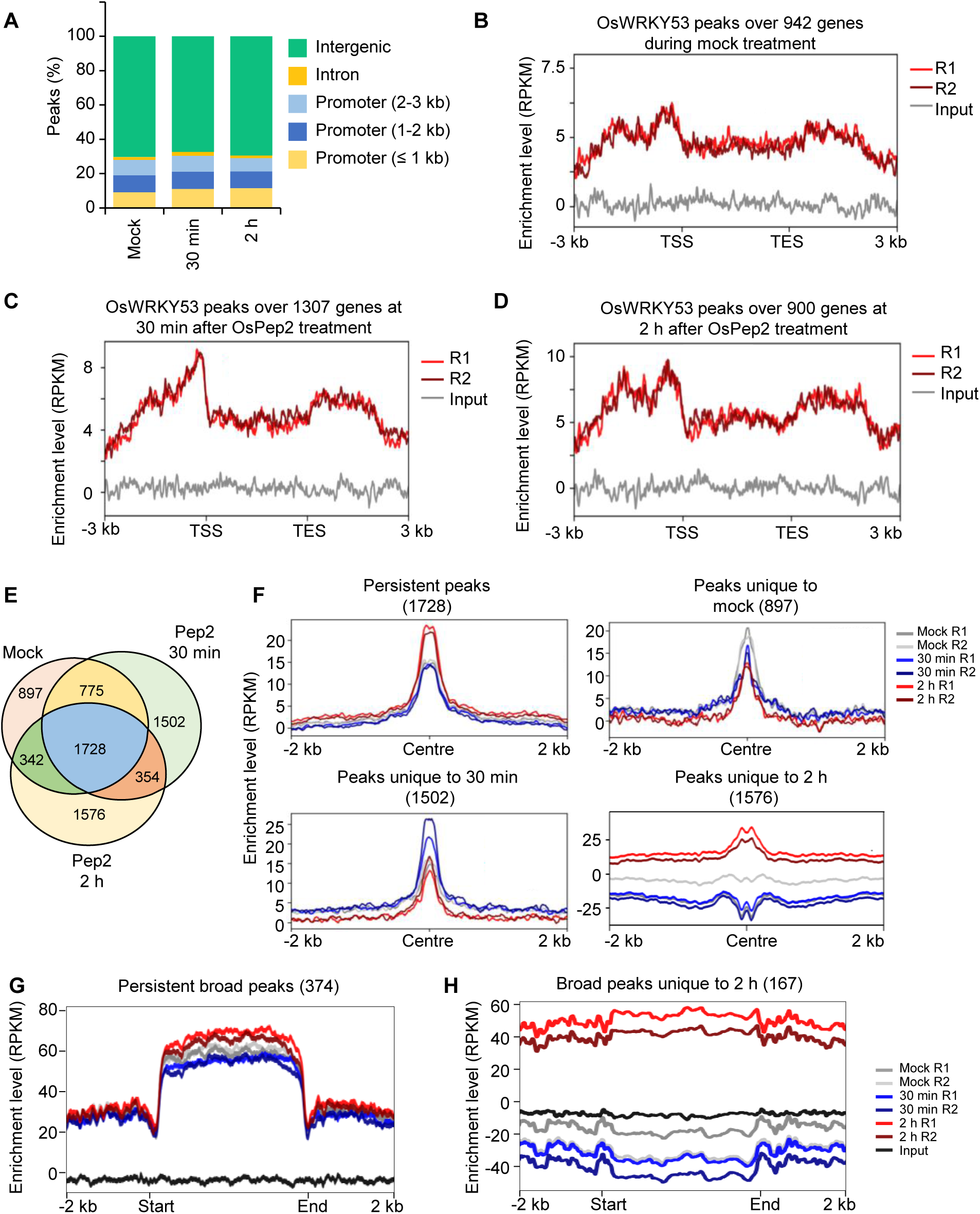
OsWRKY53 responds to OsPep2 treatment and shows altered occupancy over chromatin. **(A)** Depiction of the distribution of OsWRKY53 peaks over different regions of the genome in response to OsPep2 treatment over time. **(B-D)** Metaplots showing the enrichment of OsWRKY53 peaks over genes during mock, 30 min and 2 h post OsPep2-treatment, respectively. **(E)** Peaks intersection of OsWRKY53 across time points upon OsPep2 treatment. **(F)** Metaplots showing the enrichment of peaks common to all the time points and unique to each time point post OsPep2 treatment. **(G)** Metaplots depicting the enrichment of long peaks persistently occupied by OsWRKY53 across all time points post OsPep2-treatment. **(H)** Metaplots showing the long peaks occupied by OsWRKY53 uniquely at 2 h post OsPep2-treatment. R1 and R2 are replicates. Input represents the DNA before antibody-based enrichment. TSS – transcription start site, TES – transcription end site, Centre-Centre of each peak.

In order to investigate the nature of broad OsWRKY53-enriched peaks, we overlapped these regions with RNA-seq and found that there was RNA expression observed only in the 5’ end of the regions of the persistent broad peaks (Fig. S4B). This RNA expression is clearly under the positive control of OsWRKY53 as the RNA expression level reduced significantly in the KO1 plants (Fig. S4C). Further, among the broad peaks unique to 2 h post OsPep2 treatment, unlike persistent broad peaks, the expression was reduced at 2 h post treatment throughout the 167 genomic regions longer than 500 nt (Fig. S4D). Interestingly, in the KO1 plants, this expression pattern was not observed in the OsWRKY53-bound regions, while, there was upregulated expression at ∼1.5 kb upstream of the broad peaks. This unusual increase was observed in KO1 plants irrespective of OsPep2 treatment (Fig. S4E). These observations indicate that OsWRKY53 is involved in the suppression of RNA expression in the regions of long peaks or in the vicinity of them.

Further, to gain more insight, we intersected the long peaks with various genomic features and observed that the broad peaks emerged from RNA transposons predominantly, almost 75% of the 167 peaks (Fig. S4E). Increased transcription of potential non coding RNAs was also observed at the repeat regions occupied by OsWRKY53 (Fig. S4F-G). Some studies indicate the association between non coding RNAs and WRKYs and their downstream processes, and it will be interesting to study if OsWRKY53 mediated similar regulation (Zhang et al. 2016).

### OsWRKY53 positively dictates the expression of OsPep2-responsive genes

We considered all the OsPep2 responsive genes (upregulated, Log_2_FC ≥ 1) at different time points and observed that the expression of OsPep2-responsive genes was severely reduced in both the KO plants (Fig. 5A-D, Table S5). This indicates the global regulation of gene expression by OsWRKY53 in response to OsPep2 treatment. We further intersected the OsWRKY53 peaks with both upregulated as well as downregulated genes. At 30 min time point, a total of 198 peaks overlapped with upregulated genes upon OsPep2 treatment (Fig. 5E), while this number reduced to 82 genes at 2 h time point (Fig. 5E). OsWRKY53 showed sharp enrichment over the promoter regions of the 198 upregulated genes just before the TSS (Fig. 5F). The enrichment of OsWRKY53 over the promoters of 82 upregulated genes at 2 h were broad and bimodal as described above (Fig. 5G).

**Figure 5:**
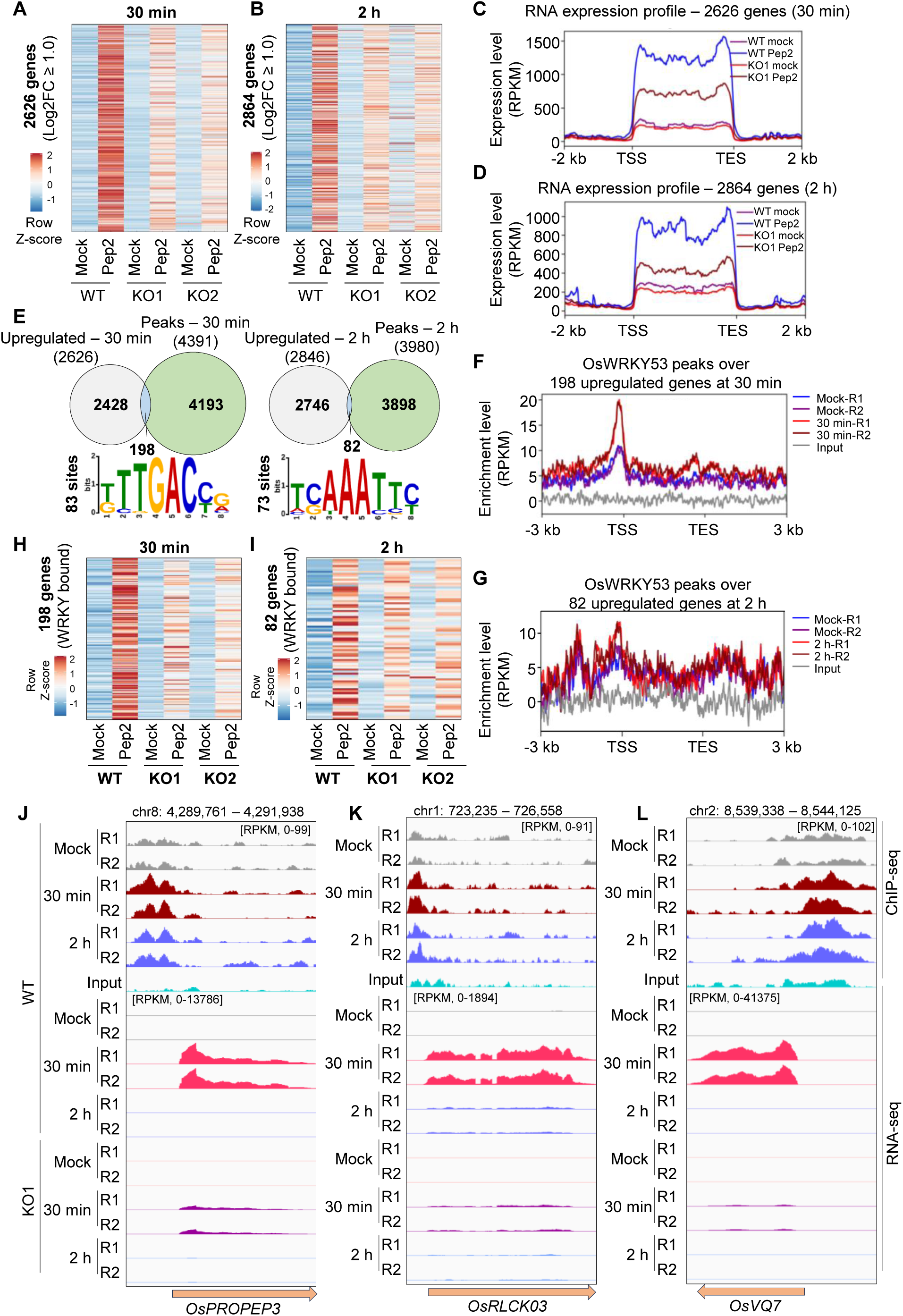
OsWRKY53 positively dictates the expression of OsPep2-responsive genes. **(A)** Heatmaps showing the RNA expression of OsPep2 responsive genes in KO and WT leaves upon OsPep2 treatment at 30 min. **(B)** Heatmaps showing the RNA expression of OsPep2 responsive genes in KO and WT leaves upon OsPep2 treatment at 2 h. **(C** and **D)** Metaplot showing the RNA expression of OsPep2 responsive genes in KO1 and WT leaves upon OsPep2 treatment at 30 min and 2 h respectively. **(E)** Overlap between genic regions of upregulated genes upon OsPep2 treatment at 30 min and 2 h. The sequence logos depict the highly enriched motifs in the promoters of the OsWRKY53 bound genes. **(F** and **G)** Metaplots showing the promoter occupancy of OsWRKY53 over upregulated genes upon OsPep2 treatment at 30 min and 2 h respectively. **(H** and **I)** Heatmaps showing the expression of OsWRKY53 bound genes upon OsPep2 treatment at 30 min and 2 h respectively. **(J**-**L)** IGV screenshots showing the expression of representative genes (*OsPROPEP3*, *OsRLCK03*, *OsVQ7*) whose promoters are occupied by OsWRKY53 in response to OsPep2 treatment.

To ascertain the nature of cis-regulatory elements involved in OsWRKY53-mediated regulation, we performed motif enrichment analysis using MEME motif tool (Bailey et al. 2015), and observed that, conventional W-box motif with a sequence of 5’- (T/G)TTGACC(G/A)-3’ (n=83) was predominantly enriched at the 30 min time point among the 198 peaks (Fig. 5E). Interestingly, at the 2 h time point though, the sequence signature was 5’-TC/GAAATTC/T-3’ (n=73), unlike a W-box motif, indicating OsWRKY53 potentially binds at different regions over time in response to OsPep2 (Fig. 5E). This altered enrichment of the motif at 2 h time point might be direct or indirect, since elaborate studies using ChIP-seq over time points during PTI have not been carried out for WRKY TFs. In spite of this, it was clear that the direct binding of OsWRKY53 to the observed motifs positively contributed to the expression of those genes as they did not respond like that of WT in both the KO lines post OsPep2 treatment (Fig. 5H-I and Fig. S5A-B, Table S6). The intersection of OsWRKY53 peaks with that of downregulated genes was poor with only 86 peaks at 30 min (Fig. S5C), possibly due to reduced or no binding over their promoters (Fig. S5D). However, as seen in KO derived RNA-seq datasets, OsWRKY53 is involved in the downregulation of substantial number of genes post OsPep2 treatment (Fig. S5E, Table S7). Over selected genes, OsWRKY53 contributed to induction of gene expression predominantly at 30 min (Fig. 5J-L). Some of the prominent genes whose expression appears to be directly regulated by OsWRKY53 include OsPROPEP3, OsRLCK03, OsVQ7, OsMYB52, OsRLCK311 and OsJMJ706 (Fig. 5J-L and Fig.S5F-H) having well-established functions in wound responses, nitric oxide signalling, salinity tolerance, and floral organ development (Xixu et al. 2020; Harshith et al. 2024; Sade et al. 2020; Sun & Zhou 2008). These results together suggest that OsWRKY53 is a master TF that contributes to a major share of gene expression changes observed upon OsPep2 treatment.

### OsWRKY53 dictates the expression of other WRKY TFs both positively and negatively

WRKY group of TFs have been predominantly shown to be substrates of MPKs. In order to check if OsWRKY53 undergoes phosphorylation upon OsPep2 perception to potentially alter binding abilities, we treated rice leaf strips with OsPep2. We observed that OsWRKY53 accumulates both in phosphorylated as well as non-phosphorylated forms at the mock treatment conditions. Upon OsPep2 peptide treatment, but not with PSK treated control, we observed specific increase in the phosphorylated form of OsWRKY53 specifically at 30 min post-treatment (Fig. 6A). This data suggests that binding of specific motifs requires phosphorylation of OsWRKY53 immediately after signal perception similar to other WRKYs studied (Zhou et al. 2020).

**Figure 6:**
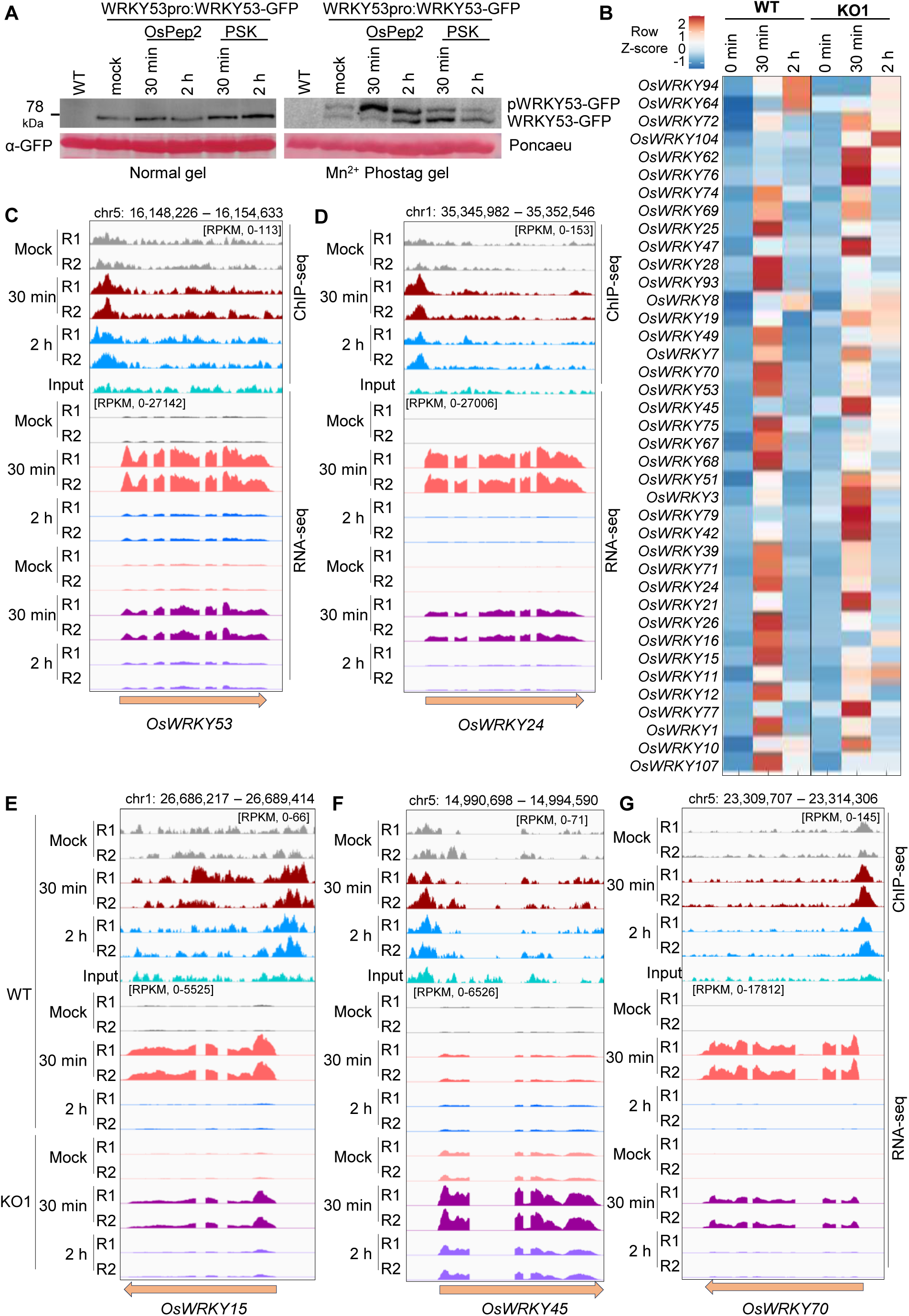
OsWRKY53 dictates the expression of WRKY TFs both positively and negatively. **(A)** Western blot images showing the response of OsWRKY53-GFP to OsPep2 and PSK treatment. The left side gel is normal SDS-PAGE based and the right side gel is Mn^2+^ phos-tag gel. pWRKY53-GFP indicates the phosphorylated form of the protein. (B) Heatmaps showing the expression of 39 OsPep2-responsive WRKYs in WT as well as in KO1 plants upon OsPep2 treatment. (C) IGV screenshot showing the autoregulatory nature of OsWRKY53 where it binds to its own promoter to regulate its expression in response to OsPep2 treatment. (D-G) IGV screenshots showing the expression of other WRKYs (*OsWRKY24*, *OsWRKY15*, *OsWRKY45* and *OsWRKY70*) which are regulated by direct binding of OsWRKY53 to their promoters.

Since we observed direct binding of OsWRKY53 to the promoters of only a subset of genes, it is possible that OsWRKY53 indirectly regulates other TFs that might contribute to the gene expression changes upon OsPep2 perception. Through RNA-seq analysis, a set of 39 WRKY TFs that were showing transcriptional induction upon OsPep2 treatment were identified (Fig. 6B). Among these, a majority of the WRKYs were under positive regulation of OsWRKY53 as observed in case of KO plants where these genes were unresponsive (Fig. 6B, Table S8). However, there were also a few WRKYs that were negatively regulated (for example, OsWRKY62, 76, 47, 45, 51, 03, 79, 42, 21, 77 and 10). Similar to other WRKYs (Birkenbihl et al. 2018), OsWRKY53 positively contributes to its own induced expression upon OsPep2 perception indicating its autoregulatory nature (Fig. 6C). We also observed that OsWRKY24 and OsWRKY70, both paralogs of WRKY53, showed induced expression upon OsPep2 treatment and this induction was dependent on OsWRKY53 binding to their promoter upon OsPep2 perception (Fig. 6D and 6G). The other positively regulated WRKYs included OsWRKY15 and OsWRKY93 (Fig. 6E and Fig. S6A), among which WRKY93 was previously implicated in the regulation of leaf senescence and biotic stress response (Li et al. 2021).

OsWRKY53 also directly bound to the promoter of OsWRKY45, a WRKY involved in both biotic and abiotic stress responses, and led to its suppression, (Shimono et al. 2012; Tao et al. 2011). In the absence of OsWRKY53, the expression of OsWRKY45 increased drastically in comparison to WT plants (Fig. 6F). OsWRKY42 is also negatively regulated by OsWRKY53 by directly binding to its promoter (Fig. S6B). Further, we also observed that the induction of some WRKYs is independent of OsWRKY53 (Fig. S6C). These results collectively indicate that OsWRKY53 is a master TF in regulating gene expression upon OsPep2 perception. OsWRKY53 cross-regulated the expression of multiple other WRKYs directly or indirectly as proposed for several WRKY TFs (Birkenbihl et al. 2018) and it will be interesting to study how these regulations are achieved and how these alterations mediate specific processes.

### OsWRKY53 temporally regulates early and late wound-responsive genes

To explore the contribution of OsWRKY53 to wound responsive gene expression, we performed wounding experiments in KO1 line and compared its transcriptome with that of WT across early, intermediary and late time points. We observed positive regulation of wound responsive genes dictated by OsWRKY53 at early time points, as both upregulation and downregulation of genes were clearly reduced in KO1 (Fig. 7A, Table S9). To ascertain the differences between the role of OsWRKY53 in wounding vs OsPep2 signalling at different time points, we categorized the common genes as well as unique genes responding to either wounding or OsPep2 treatment across both the time points (Fig. 7B). OsWRKY53 positively contributed to induced expression of wound as well as OsPep2-specific genes at 30 min (Fig. 7C, Table S10). However, at 2 h, wound-specific genes were negatively regulated, while OsPep2-responsive genes were positively regulated by OsWRKY53, indicating its differential contribution to wound and OsPep2 signalling (Fig. 7C). Very interestingly, in the absence of OsWRKY53, both the wound induced upregulation and downregulation of genes were completely suppressed at 12 h time point (Fig. 7A). This indicated that the late wound responses in Oswrky53 KO plants phenocopied the suppressed wound responses displayed by OsPSKR OE lines in terms of transcriptome that we previously described (Harshith et al. 2024). The set of 53 early wound responsive genes which were directly downstream to OsPSKR (Harshith et al. 2024) were suppressed in KO1 plants upon wounding, conclusively indicating that OsWRKY53 is involved in the regulation of wound-responsive genes downstream of OsPSKR signalling (Fig. 7D, Table S11). These results collectively suggest that OsWRKY53 is involved in temporal fine-tuning of gene expression upon wounding.

**Figure 7:**
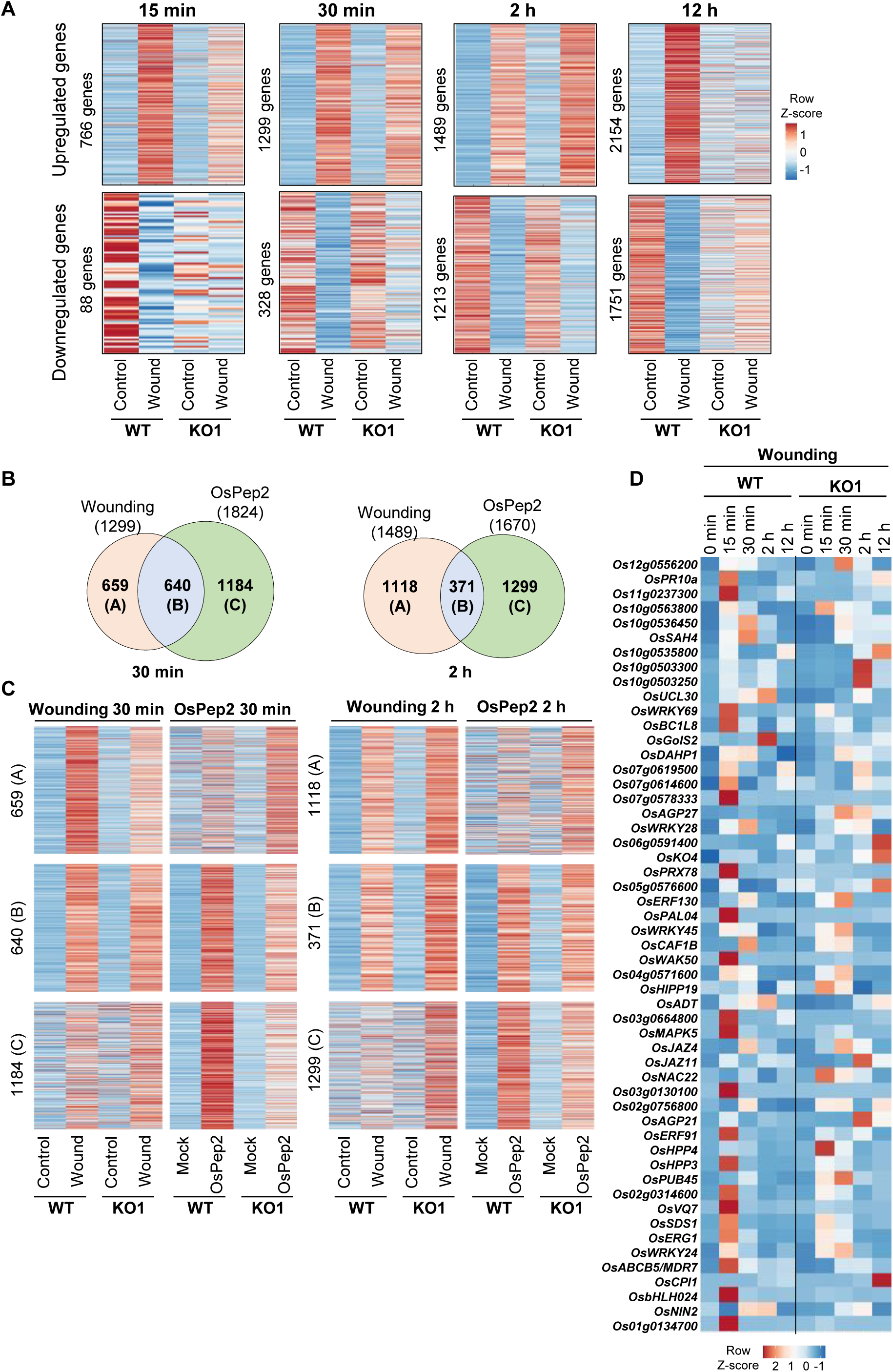
OsWRKY53 dictates wound-responsive genes distinctly at early time point in comparison to late time point. **(A)** Heatmaps showing the expression of wound-responsive genes across time upon wounding in KO1 plants compared to WT plants. **(B)** Overlap between wound-responsive and OsPep2-responsive genes at 30 min and 2 h post treatment/injury. **(C)** Heatmaps showing the category-wise (corresponding to **(B)**) expression pattern of wound-responsive and OsPep2-responsive genes at 30 min and 2 h post treatment/injury. **(D)** Heatmap showing the expression pattern of 53 wound-responsive genes that are directly downstream to OsPSKR.

### PSK-responsive genes are constitutively mis-regulated in the absence of OsWRKY53

Since, Oswrky53 KO plants phenocopied the transcriptome of OsPSKR OE, we hypothesized that OsWRKY53 and OsPSKR play contrasting signalling roles. In order to obtain a detailed understanding of this cross-talk, we compared the mis-regulated genes between Oswrky53 KO and OsPSKR OE. We observed a huge overlap between these two gene sets indicating that indeed signalling dictated by OsPSKR and OsWRKY53 are in contrast to each other (Fig. 8A). The genes upregulated in OsPSKR OE and Oswrky53 KO1 were predominantly involved in photosynthesis (Fig. S7A). This suggests the positive role of OsPSKR in growth and the negative role of OsWRKY53 in dictating growth responses. The downregulated genes in OsPSKR OE and Oswrky53 KO were mainly involved in rRNA processing and translation related processes (Fig. S7B).

**Figure 8:**
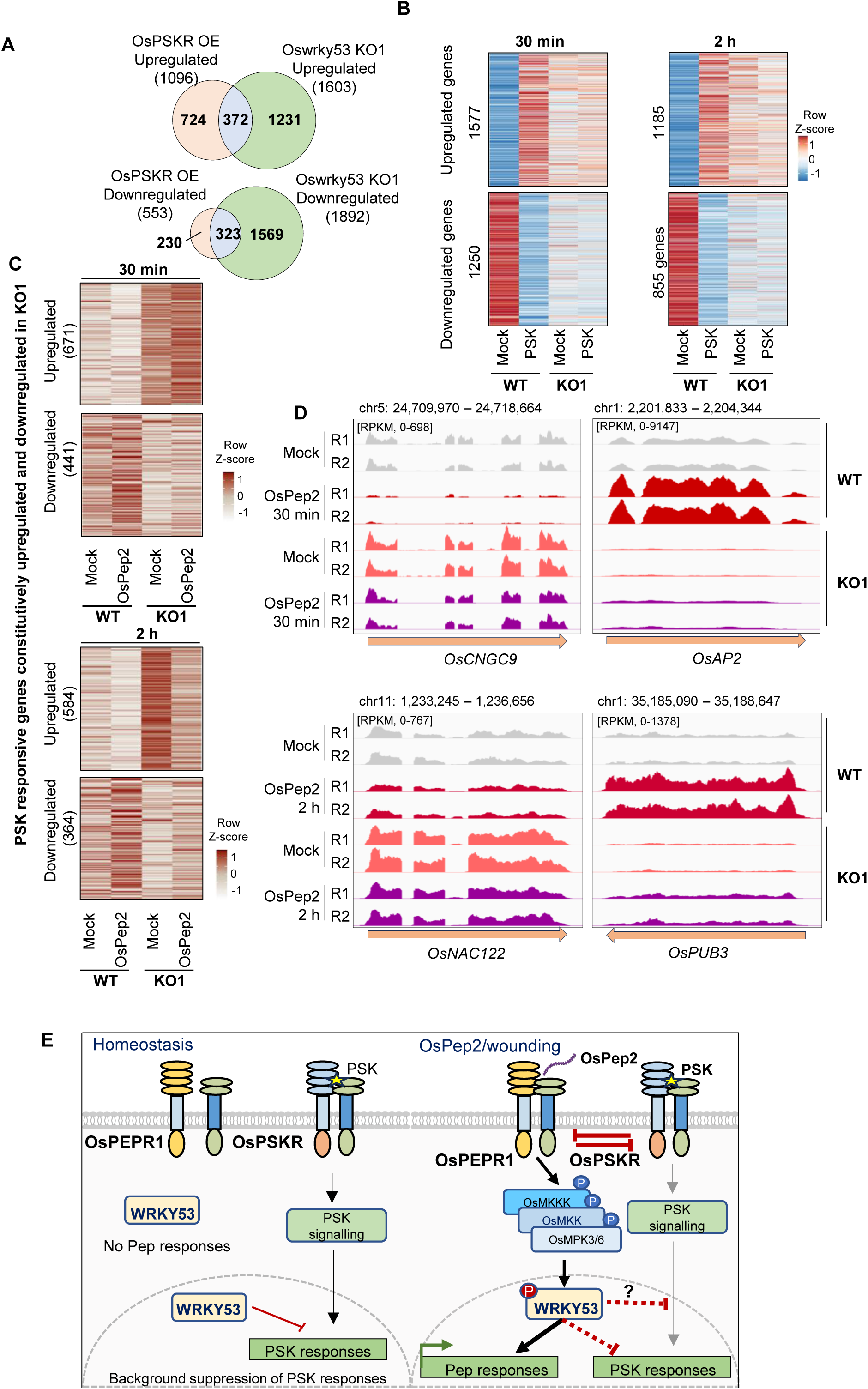
PSK-responsive genes are constitutively mis-regulated in the absence of OsWRKY53. **(A)** Overlap of genes mis-expressed in OsPSKR OE and Oswrky53 KO plants. **(B)** Heatmaps showing the expression pattern of PSK-responsive genes in WT and KO1 plants in response to PSK treatment at 30 min and 2 h. **(C)** Heatmaps showing the expression pattern of PSK-responsive genes that were constitutively perturbed (Log_2_FC ≥ 1.5 for upregulated and Log_2_FC ≤ -1.5 for downregulated) in KO1 in comparison to WT during homeostasis in response to OsPep2 treatment at 30 min and 2 h. **(D)** IGV screenshots depicting the expression pattern of representative PSK responsive genes that are cross-regulated by OsWRKY53 during response to OsPep2 perception. **(E)** Model showing the role of OsWRKY53 in regulation of gene expression in response to wounding/OsPep2/PSK treatment. During homeostasis there is basal level expression of OsWRKY53 that predominantly exists in non-phosphorylated form. In the absence of defense signal activation, OsWRKY53 contributes to brassinosteroid signalling among other metabolic pathways. Also, OsWRKY53 suppresses background expression of PSK responsive genes. Upon defense signal activation through wounding or OsPep2 treatment, OsWRKY53 expression gets induced. Further, OsWRKY53 undergoes phosphorylation specifically upon OsPep2 perception. This positively contributes to gene expression downstream to OsPep2 signalling. OsWRKY53 simultaneously suppresses PSK-responsive genes when OsPep2 signalling is turned on. Together, OsWRKY53 is predominantly involved in positive regulation of early defense responses. It contributes to fine tuning of cross-talk involved between defense and growth responses, especially between OsPep2 and PSK mediated signalling.

Further, to check the role of OsWRKY53 in mediating gene expression upon PSK perception, we treated Oswrky53 KO1 plants with PSK and performed transcriptome profiling at 30 min and 2 h post treatment. The results obtained indicated that PSK responsive genes were constitutively differentially expressed in KO1 plants and the response to PSK treatment was compromised in KO1 plants (Fig. 8B, Table S12). Since OsWRKY53 is a positive regulator of responses downstream to OsPep2 treatment, we explored if PSK responsive genes that are constitutively perturbed in KO1 are involved in OsPep2 responses. Interestingly, both during 30 min and 2 h, PSK responsive genes in WT conditions, but not in Oswrky53 KO1, were suppressed upon treatment of plants with OsPep2 (Fig. 8C-D, Table S13). These results suggest that OsWRKY53 acts as the gatekeeper of PSK responsive genes during homeostasis as its absence leads to constitutive upregulation of PSK-responsive genes. During OsPep2 perception, OsWRKY53 is involved in positive regulation of gene expression changes downstream to OsPep2 pathway while the PSK-responsive genes are simultaneously suppressed. In the absence of OsWRKY53, this cross-regulation of the expression of PSK-responsive genes during activation of OsPep2 signalling is lost (Fig. 8E). All these results collectively suggest that OsWRKY53 is very important for fine-tuning of wound-responses through modulation of both PEP and PSK-responsive genes and acts at the intersection of growth vs defense.

## Discussion

The waves of transcription leading to expression and suppression of distinct categories of genes dictate fine-tuning of physiological and phenotypic outputs across plants. The ultimate outcome of a wounding event in dicots is wound-induced regeneration/growth (Ikeuchi et al. 2017). This outcome is a result of interplay between major signalling events primarily mediated by specific family of TFs, majorly belonging to AP2 domain containing family (Ikeuchi et al. 2020). Monocots, however, do not possess this regeneration ability (Liang et al. 2023). Nevertheless, wounding in monocots promotes elongation of leaf blades, promotes growth of new shoots and restricts wound-induced cell death. This necessitates a tightly coordinated transcriptional regulation primarily dictated by specific TFs potentially distinct from what is observed in dicots. Investigation of the role of a specific TF in this context is challenging as many genes that code for TFs show transcriptional upregulation or downregulation upon wound or wound-derived PEP perception. Several of these TFs have functional redundancy. The major emphasis is on the TFs that can promote growth or regeneration post-wounding, often acting at the intersection of signal transitions (Zhang et al. 2022; Heyman et al. 2018; Ikeuchi et al. 2020). The TFs that promote defense while simultaneously suppressing growth responses are equally important for a successful response. There are studies in rice that show the involvement of a single TF (IPA1) in regulating both yield and immunity (Wang et al. 2018). Similar roles for WRKYs during transition of wound responses have not been demonstrated earlier. Identification of the role of OsWRKY53 in promoting defense responses while suppressing growth responses is interesting in that it seamlessly integrated and counteracted with PSK-mediated growth promotion that we identified previously (Harshith et al. 2024). These results are in agreement with the nature and functions of OsWRKY53 interacting partners, OsVQ25 and OsDLA, in mediating responses during infections (Hao et al. 2022; Meng et al. 2024). Novel insights regarding the selective transcription regulated by OsWRKY53 during activation of PTI described here might be applicable to other stresses.

Rice has over a hundred WRKYs and it has been challenging to study properties of individual WRKYs due to functional redundancy between related WRKYs (Chen et al. 2019). OsWRKY53 appears to be a predominant TF involved in regulation of gene expression changes upon OsPep2 treatment and wound responses. A sharp transcriptional response to PEP treatment at the early time point is mediated through OsWRKY53. We observed a direct regulation of OsPROPEP3 expression upon OsPep2 by OsWRKY53 through binding to W-boxes. Similar regulation of PEP coding genes by AtWRKY33, which is an ortholog of OsWRKY53, has been demonstrated suggesting the conservation of this signalling across plants (Logemann et al. 2013). While the bulk of the transcriptional changes upon OsPep2 treatment were affected in OsWRKY53 KO plants, OsWRKY53 displayed direct occupancy of the promoters of only a subset of genes. Hence it is possible that, contribution of OsWRKY53 to expression of unbound genes could be due to indirect effects. Since ChIP-seq datasets for related PTI-related WRKYs are not available in model systems *Arabidopsis* or rice, it is difficult to understand if WRKY bound regions show conservation across plants. It has been argued that OsWRKY53 acts as a negative regulator of defense responses against insect herbivory based on the performance of insect larvae (Hu et al. 2015). However, we observe it to be positively contributing to PEP induced defenses. It would be interesting to investigate if the role of OsWRKY53 in herbivory response is uncoupled from PEP responses or this depends on the host plant. In agreement with our findings here, Dhar et al. (Dhar et al. 2024), have recently identified a TF involved in growth vs defense transitions in *Arabidopsis* upon PEP-mediated signalling.

WRKY TFs act in a network leading to multiple cross-regulation of other WRKYs during immune signal activation in *Arabidopsis* (Liu et al. 2024). We also observed cross-regulation of multiple WRKYs through direct binding of OsWRKY53 to their promoters. This suggests that OsWRKY53 acts as one of the major TFs involved in regulation of early defense responses in rice. Interestingly, OsWRKY53 occupied a unique set of genomic regions at 2 h post PEP treatment. It is intriguing why OsWRKY53 relocalizes to non-genic regions over time. Rice intergenic regions are abundant with TEs, and in agreement with this, peaks emerging 2 h post OsPep2 treatment overlapped with several RNA TEs. Several non-coding RNAs responding to stresses including wounding have been identified (Huang et al. 2023), and it is possible that here again OsWRKY53 regulates such RNAs to mediate specific processes. However, the DEGs 2 h post wounding were mostly enriched with rRNA processing, ncRNA processing and regulation of translation. Additionally, we also observed that the CREs present in the OsWRKY53-bound regions changed over time post PEP treatment, and interestingly the regions of more than 2 kb were enriched unlike at early time points. In such peaks, WRKY bound regions spread out encompassing regions potentially generating ncRNAs. At least two examples of alterations of expression of RNAs in these regions were observed in OsWRKY53 dependent manner (Fig. S4F-G). Also, a cross-talk has been observed between AtWRKY53 and a histone deacetylase, AtHDA9 in regulating stress responses in *Arabidopsis* (Zheng et al. 2020). It will be interesting to study the functions of associated epigenetic regulation of WRKYs in detail.

Many reports have suggested the possibility of binding of WRKYs to CREs other than W-boxes (Birkenbihl et al. 2018). It is also possible that the enrichment of CREs observed at 2 h post OsPep2 are indirect through other interacting TFs or DNA binding proteins. We detected that OsWRKY53 undergoes phosphorylation at early time point post OsPep2 treatment. Previous studies have shown that OsWRKY53 can act as substrates to at least two categories of kinases, i.e., MPKs and GSKs, leading to distinct fates (Tian et al. 2021). *Arabidopsis* homolog of OsWRKY53, AtWRKY33, undergoes differential phosphorylation by CPKs and MPKs (Zhou et al. 2020). There is also cross-talk with SUMOylation of AtWRKY33 (Verma et al. 2021), suggesting multiple PTMs and their interplay might dictate the function of these TFs. Interestingly, there is a debate if phosphorylated or non-phosphorylated OsWRKY53 localizes to nucleus to mediate gene expression *via* binding to W-boxes (Meng et al. 2024). Since at 30 min post OsPep2 treatment OsWRKY53 underwent phosphorylation (Fig. 6A), we performed ChIP-seq in this time point, and observed clear binding to W-boxes. It is also possible that the differences in binding of OsWRKY53 to different regions at later time point (2 h) might be due to altered preference of non-phosphorylated OsWRKY53. It will be interesting to study binding preferences of phospho-mimics of OsWRKY53 under OsPep2 perception, however, as expected among different WRKY TFs, there are several potential phosphorylation sites in OsWRKY53.

Wound responses are dynamic and this necessitates the differential regulation of signalling. We observed that OsWRKY53 is a positive regulator of early wound responses and a negative regulator of genes expression 2 h post wounding. Strikingly, the gene expression status at 12 h time post-wounding in OsWRKY53 KO plants were similar to the gene expression status at 12 h post-wounding in OsPSKR OE that we observed in our previous work (Harshith et al. 2024). This indicates that OsWRKY53 potentially antagonizes gene expression downstream of PSK signalling. Corroborating this idea, the PSK responsive genes were constitutively mis-regulated in Oswrky53 KO plants. Also, the PSK responsive genes were actively suppressed by OsWRKY53 when OsPep2 responses were active. Such a relationship for a WRKY TF and a growth promoting module such as PSK:PSKR has not been documented earlier. Collectively, our experiments indicate that OsWRKY53 is a very important TF in regulation of wound as well as OsPep2 responses where it positively regulates early responses and assists in controlled expression of PSK responsive genes. Further studies investigating the protein interaction dynamics of OsWRKY53 in response to wounding, PEP treatment and PSK treatment would shed light on the mechanistic basis of wound responsive signalling regulation mediated by OsWRKY53. Since homologs of OsWRKY53 are found across monocots having distinct wound responsive strategies, it will be interesting to study if those homologs play similar roles during wound-related stresses.

## Acknowledgements

We thank all the members of Shivaprasad lab for reading through the manuscript and inputs. We thank the Next Generation Genomics and radiation facilities at NCBS-TIFR, Bangalore. We extend thanks to K. Veluthambi for binary vectors, PB1 seeds and *Agrobacterium* strains. This study was supported by Department of Atomic Energy, Government of India, under Project Identification No. RTI 4006 (1303/3/2019/R&D-II/DAE/4749 dated 16.7.2020).

## Author contributions

C.Y.H. and P.V.S. conceptualized the study and wrote the manuscript. C.Y.H. performed most of the experiments and analysed the data. R.D. helped in generating KO transgenic plants. D.P. helped with protein work.

## Data availability

All data generated or analysed during this study are included in this published article (and its supplementary information files). RNA-Seq data is available under GEO accession numbers; GSE260646 and GSExxxxxx. ChIP-seq data is available under GEO accession number; GSExxxxxx.

## Supplementary data

This manuscript has 7 supplementary figures and 14 supplementary tables.

**Figure S1: Expression analysis of wound as well as PEP-responsive genes that are downstream to OsPSKR**

**Figure S2: Generation and analysis of OsWRKY53 transgenic lines driven by its native promoter**

**Figure S3: OsWRKY53 occupies distinct genomic regions at 2 h upon OsPep2 treatment**

**Figure S4: OsWRKY53 occupies broader region and dictates RNA expression in the region**

**Figure S5: OsWRKY53 does not directly occupy the promoters of downregulated genes upon OsPep2 treatment.**

**Figure S6: OsWRKY53 regulates the expression of other WRKYs**

**Figure S7: Similar categories of genes are contrastingly regulated by OsPSKR and OsWRKY53**

