## Supplementary material for "OsWRKY53 dictates wound responses in rice through fine-tuning cross-talk between PEP and PSK mediated signalling": Table S14

**Table S14: List of primers used in this study**

| **OsWRKY53_qF** | CTTCGTCACGGTCTCCTCGCTTC |
| --- | --- |
| **OsWRKY53_qR** | GCTGCGCGGACTTGAACTTG |
| **OsActin_qF** | GCTATGTACGTCGCCATCCAGG |
| **OsActin_qR** | TGAGATCACGCCCAGCAAGG |
| **OsWRKY53_gRNA1_pRGEB1_R** | AATTAGGTCTCAAAACCTGAGCCGGGATTGCACCGGTGCCACGGATCATCTGCACAAC |
| **OsWRKY53_gRNA2_pRGEB1_R** | AATTAGGTCTCAAAACCGTCGAGCAACAAGAGCGGCTGCCACGGATCATCTGCACAAC |
| **wrky53 KO guide RNA1** | CCGGTGCAATCCCGGCTCAG |
| **wrky53 KO guide RNA2** | GCCGCTCTTGTTGCTCGACG |
| **OsWRKY53_pro_F_BamHI** | CTGCATGGATCCCCTCCAGATCACTTGAC |
| **OsWRKY53_pro_R_SalI** | ACTATCGTCGACGGCGAGCGTACCACAGC |
| **OsWRKY53_CDS_F** | GGTGTTACTTCTGCAGGTCGACTCTAGAGGATCCATGGCGTCCTCGACGGGG |
| **OsWRKY53_CDS_R** | GCAGAGGAGCGACTCGACGAACAGG |
| **GFP_F** | CTGTTCGTCGAGTCGCTCCTCTGCGGAATGGTGAGCAAGGGCGAGGAGCTGTTC |
| **GFP_R** | ATTTCAGCGTACCGAATTCGAGCTCGTTACTTGTACAGCTCGTCCATGCCGAGAG |
