## Supplementary Figures for "OsWRKY53 dictates wound responses in rice through fine-tuning cross-talk between PEP and PSK mediated signalling"

### Supplementary information

Manuscript has 7 Supplemental Figures and 14 Supplemental Tables.

### Supplementary figures:

**Figure S1:** Expression analysis of wound as well as PEP-responsive genes that are downstream to OsPSKR

**Figure S2:** Generation and analysis of OsWRKY53 transgenic lines driven by its native promoter

**Figure S3:** OsWRKY53 occupies distinct genomic regions at 2 h upon OsPep2 treatment

**Figure S4:** OsWRKY53 occupies broader region and dictates RNA expression in the region

**Figure S5:** OsWRKY53 does not directly occupy the promoters of downregulated genes upon OsPep2 treatment.

**Figure S6:** OsWRKY53 regulates the expression of other WRKYs

**Figure S7:** Similar categories of genes are contrastingly regulated by OsPSKR and OsWRKY53

### Supplementary Tables:

**Table S1:** Row Z-score values of WRKY expression levels related Fig. 1C-D

**Table S2:** FPKM values of 39 genes related to Fig. S1B-C

**Table S3:** FPKM values of DEGs related to Fig. 2F

**Table S4:** FPKM values of genes related to Fig. S2F-H

**Table S5:** FPKM values of genes related to Fig. 5A-B

**Table S6:** FPKM values of genes related to Fig. 5H-I

**Table S7:** FPKM values of genes related to Fig. S5E

**Table S8:** FPKM values of WRKYs related to Fig. 6B

**Table S9:** FPKM values of genes related to Fig. 7A

**Table S10:** FPKM values of genes related to Fig. 7C

**Table S11:** FPKM values of genes related to Fig. 7D

**Table S12:** FPKM values of genes related to Fig. 8B

**Table S13:** FPKM values of genes related to Fig. 8C

**Table S14:** List of primers used in this study

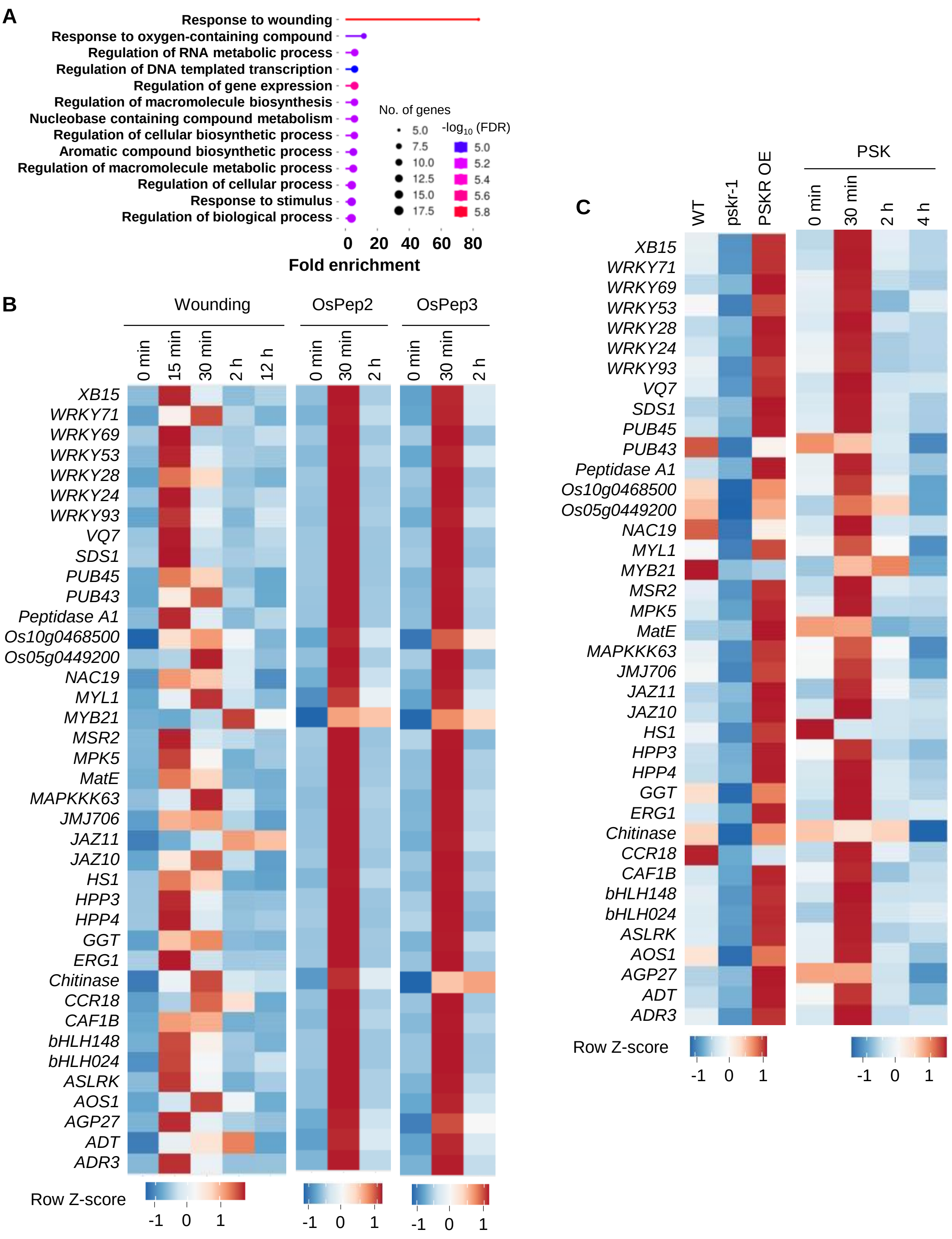


**Figure S1: Expression analysis of wound as well as PEP-responsive genes that are downstream to OsPSKR**

**(A)** GO analysis of the 39 genes downstream to OsPSKR that are wound-responsive. **(B)** Heatmaps representing the expression of 39 genes downstream to OsPSKR that are wound-responsive upon wounding and wound-derived PEP treatments. **(C)** Heatmaps representing the expression of 39 genes downstream to OsPSKR that are wound-responsive in OsPSKR mis-expression lines and upon PSK treatment. For **(B** and **C)**, gene expression is represented as row Z-score of FPKM obtained from average of replicates in RNA-seq experiments.

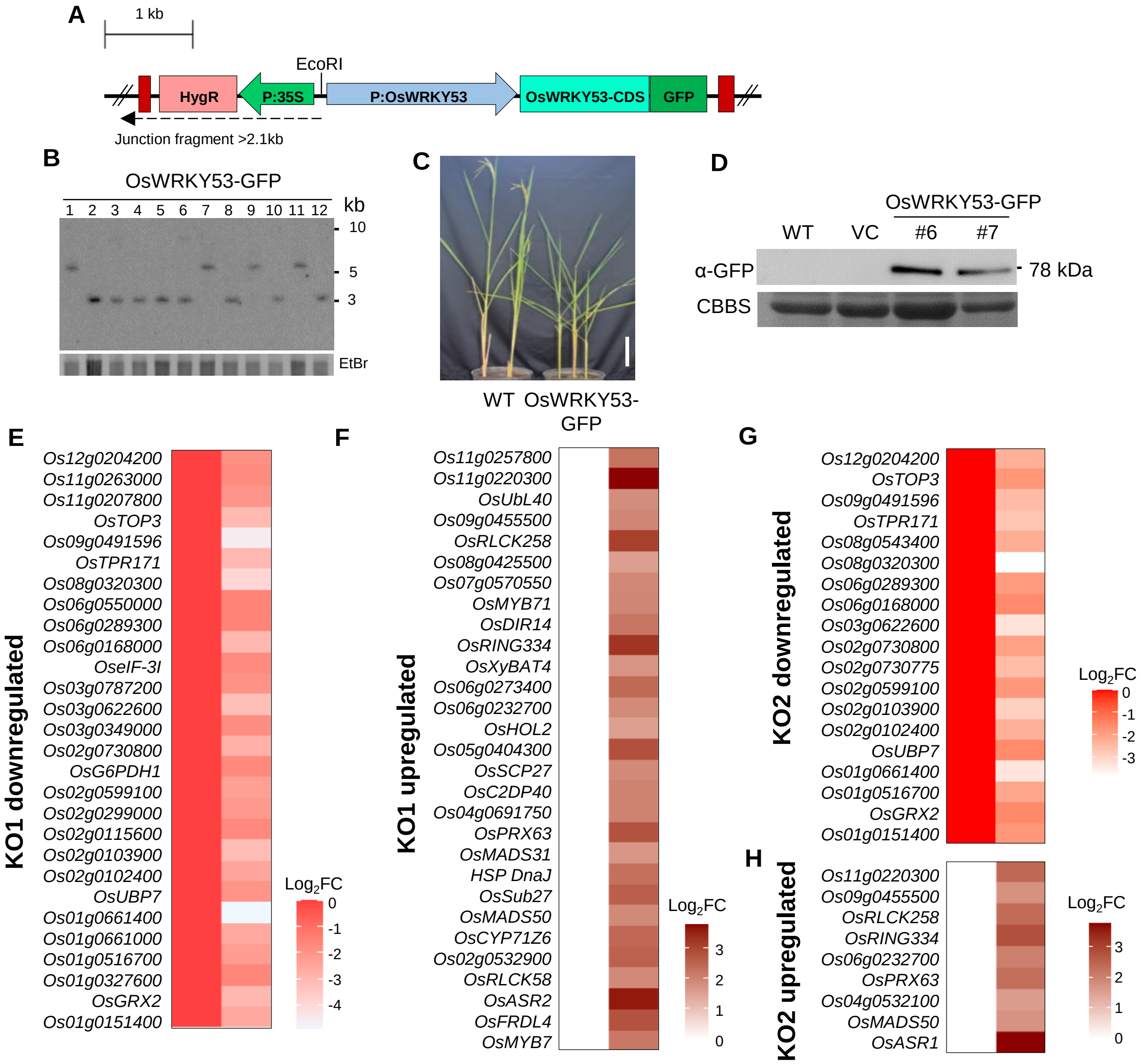


**Figure S2: Generation and analysis of OsWRKY53 transgenic lines driven by its native promoter**
**(A)** Linear map showing T-DNA region of the construct used for expressing OsWRKY53-GFP under its own promoter. **(B)** Junction fragment Southern analysis confirming the nature of transgenes. The junction fragments should be more than 2.1 kb. **(C)** Phenotypes of 14-week-old transgenic plants expressing OsWRKY53-GFP under its promoter. Scale - 8 cm. **(D)** Western blot depicting the expression of OsWRKY53-GFP in transgenic plants. **(E-H)** Expression of OsWRKY53 bound genes in KO plants compared to WT.

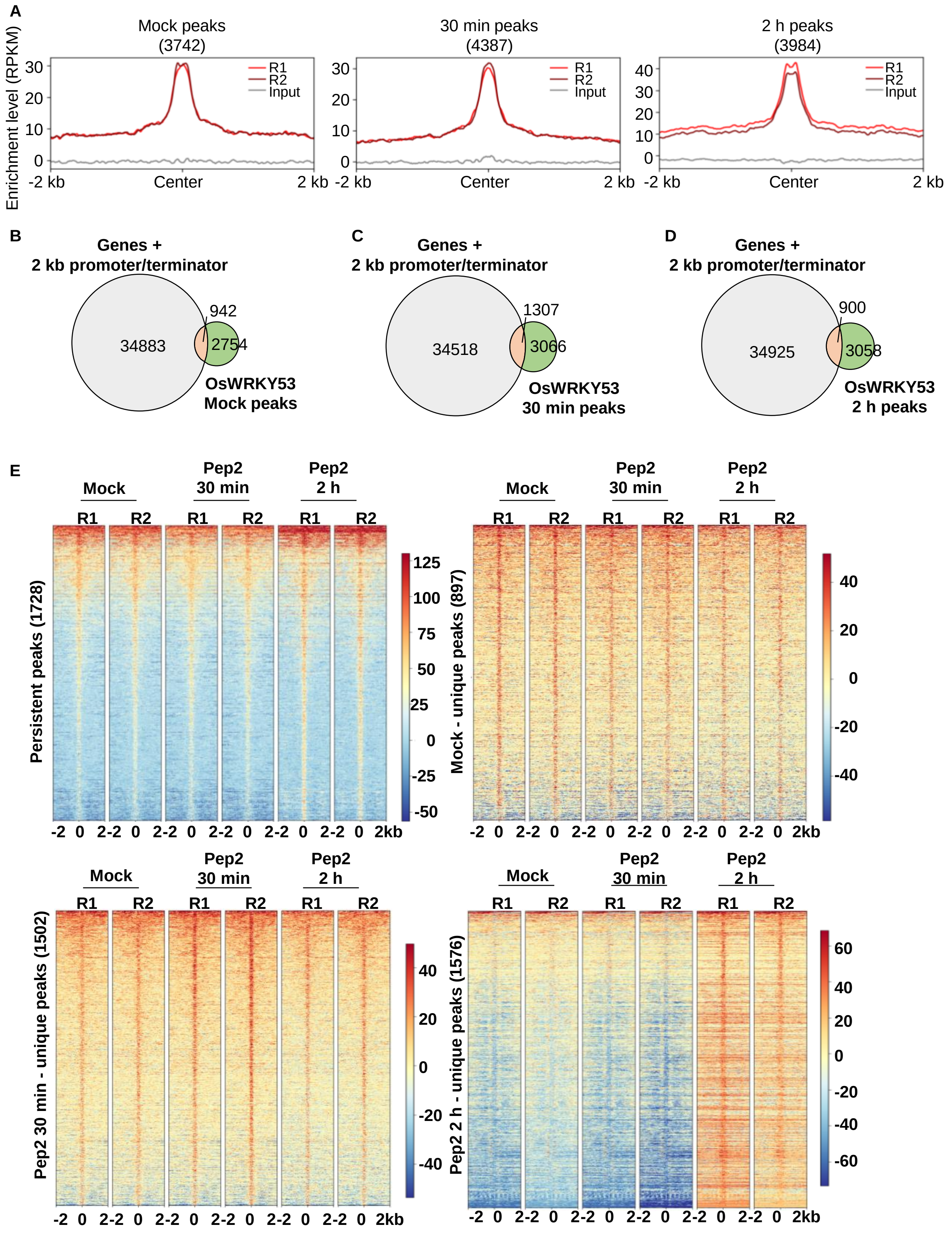


**Figure S3: OsWRKY53 occupies distinct genomic regions at 2 h upon OsPep2 treatment.**

**(A)** Metaplots showing the enrichment of OsWRKY53 peaks across time post OsPep2-treatment. **(B-D)** Overlap of OsWRKY53 peaks with the genic regions along with 2 kb promoter/terminator regions across time post OsPep2-treatment. **(E)** Heatmaps showing the enrichment of OsWRKY53 peaks over the regions persistently occupied and uniquely occupied at each time point post OsPep2-treatment. For heatmaps, enrichment level (RPKM) is shown as color coded scale.

A

| Length of peak (nt) | Mock (897) | 30 min (1502) | 2 h (1576) | Persistent (1728) |
|---|---|---|---|---|
| 0-200 | 701 | 1112 | 801 | 545 |
| 200-500 | 196 | 386 | 608 | 809 |
| 500-1000 | - | 4 | **123** | **247** |
| >1000 | - | - | **44** | **127** |

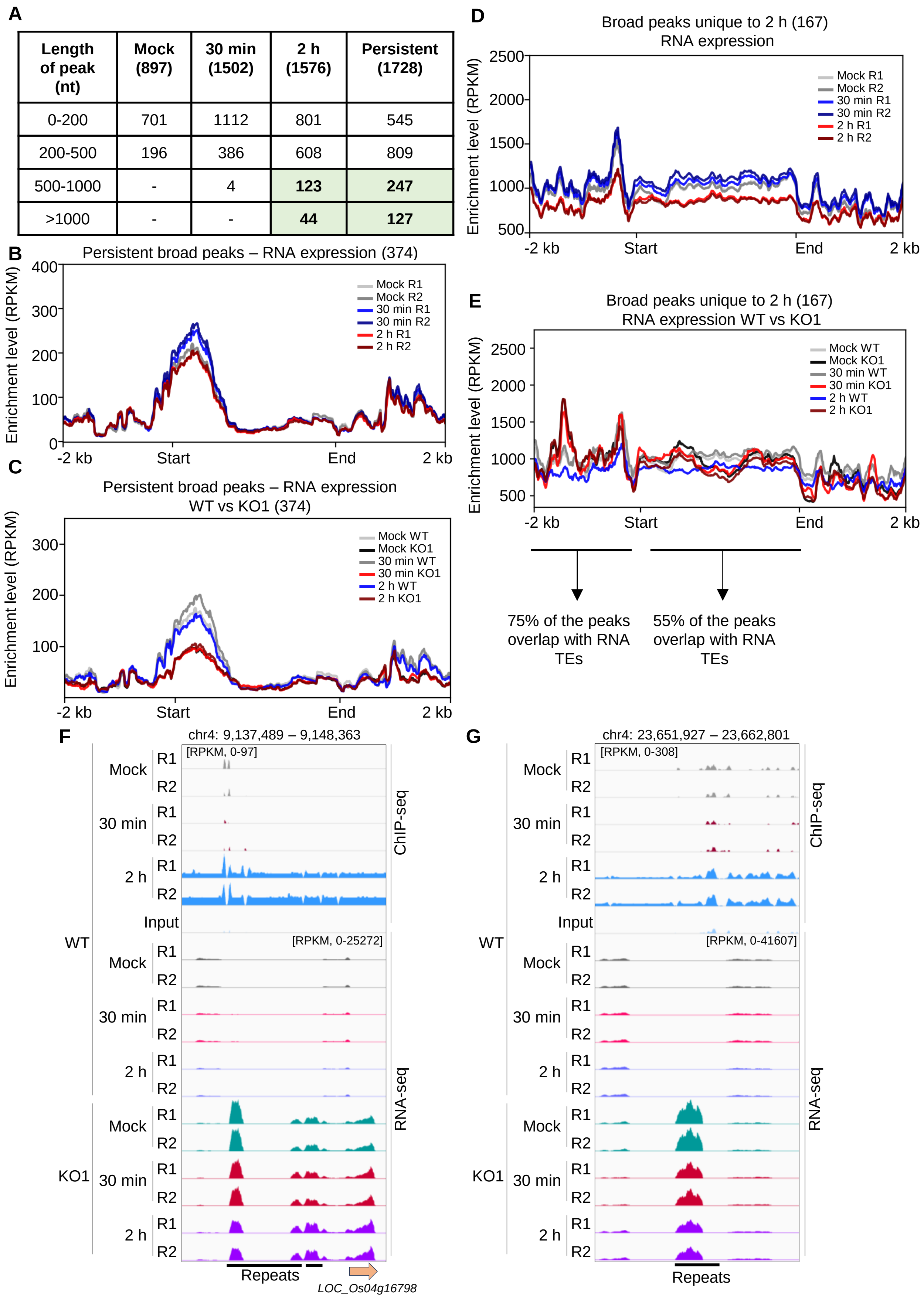

**Figure S4: OsWRKY53 occupies broader region and dictates RNA expression in the region**

**(A)** Tabular representation of the length of the peaks occupied by OsWRKY53 across time post OsPep2-treatment. (B) Metaplots showing the RNA expression profile in the broad regions occupied by OsWRKY53 persistently. (C) Metaplots showing the RNA expression profile in the broad regions occupied by OsWRKY53 persistently in KO1 plants compared to WT. (D) Metaplots showing the RNA expression profile in the broad peak regions occupied by OsWRKY53 uniquely at 2 h post OsPep2-treatment. (E) Metaplots showing the RNA expression profile in the broad peak regions occupied by OsWRKY53 uniquely at 2 h post OsPep2-treatment in KO1 plants compared to WT. The text below the figure represents the proportion of corresponding regions overlapping with RNA TEs. TEs – transposable elements. (F and G) IGV screenshots representing the occupancy of OsWRKY53 in broad regions and the corresponding RNA expression in WT vs KO1.

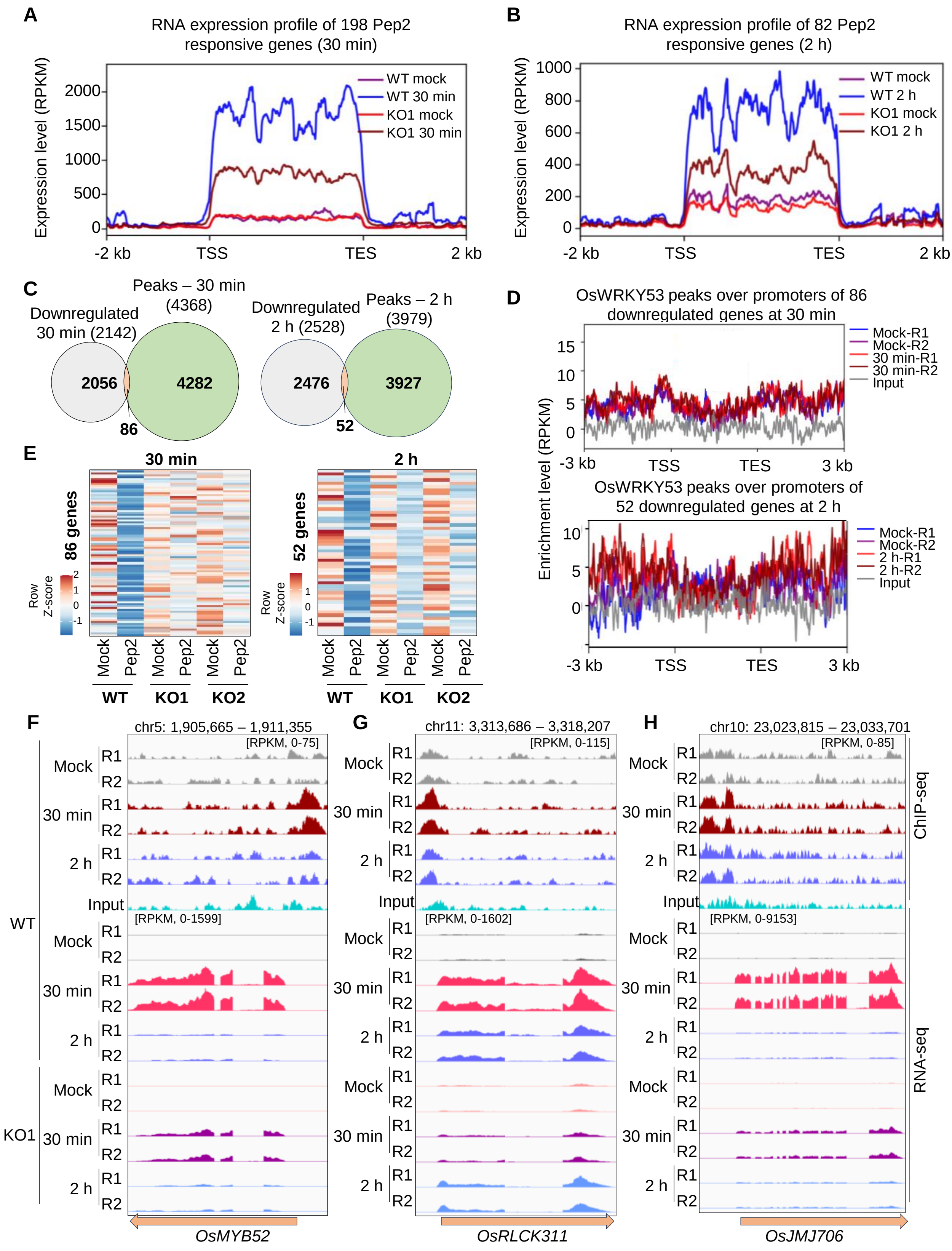
A
RNA expression profile of 198 Pep2 responsive genes (30 min)
Expression level (RPKM)
WT mock
WT 30 min
KO1 mock
KO1 30 min
-2 kb
TSS
TES
2 kb
B
RNA expression profile of 82 Pep2 responsive genes (2 h)
Expression level (RPKM)
WT mock
WT 2 h
KO1 mock
KO1 2 h
C
Downregulated 30 min (2142)
Peaks – 30 min (4368)
2056
4282
86
Downregulated 2 h (2528)
Peaks – 2 h (3979)
2476
3927
52
D
OsWRKY53 peaks over promoters of 86 downregulated genes at 30 min
Mock-R1
Mock-R2
30 min-R1
30 min-R2
Input
Enrichment level (RPKM)
-3 kb
TSS
TES
3 kb
OsWRKY53 peaks over promoters of 52 downregulated genes at 2 h
Mock-R1
Mock-R2
2 h-R1
2 h-R2
Input
E
30 min
86 genes
Row Z-score
2 h
52 genes
Mock
Pep2
WT
KO1
KO2
F
chr5: 1,905,665 – 1,911,355
[RPKM, 0-75]
[RPKM, 0-1599]
OsMYB52
G
chr11: 3,313,686 – 3,318,207
[RPKM, 0-115]
[RPKM, 0-1602]
OsRLCK311
H
chr10: 23,023,815 – 23,033,701
[RPKM, 0-85]
[RPKM, 0-9153]
OsJMJ706
Mock
30 min
2 h
Input
R1
R2
WT
KO1
ChIP-seq
RNA-seq

**Figure S5: WRKY53 does not directly occupy the promoters of downregulated genes upon OsPep2 treatment.**

**(A** and **B)** Metaplots showing the expression of genes whose promoters are directly bound by OsWRKY53 upon OsPep2 treatment at 30 min and 2 h respectively in WT and KO1 plants. **(C)** Overlap between genic regions of downregulated genes upon OsPep2 treatment and OsWRKY53 peaks at 30 min and 2 h post OsPep2 treatment. **(D)** Metaplot showing the occupancy of OsWRKY53 in the promoters of downregulated genes upon OsPep2 treatment at 30 min and 2 h. **(E)** Heatmap showing the expression of downregulated genes upon OsPep2 treatment at 30 min and 2 h. **(F-H)** IGV screenshots showing the expression of representative genes (*OsMYB52*, *OsRLCK311*, *OsJMJ706*) whose promoters are occupied by OsWRKY53 in response to OsPep2 treatment.

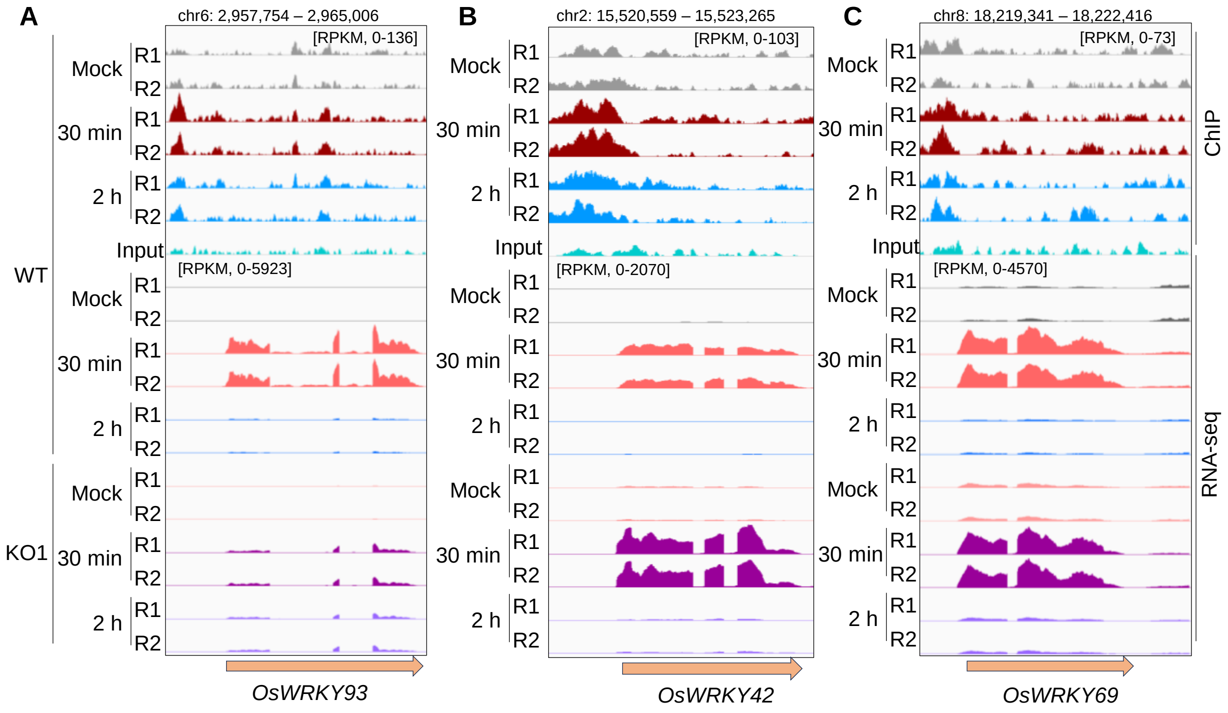


**Figure S6: OsWRKY53 regulates the expression of other WRKYs**

**(A)** IGV screenshot depicting the positive regulation of the expression of *OsWRKY93* by OsWRKY53 by directly binding to its promoter in response to OsPep2 treatment. **(B)** IGV screenshot depicting the negative regulation of the expression of *OsWRKY42* by OsWRKY53 by directly binding to its promoter in response to OsPep2 treatment. **(C)** IGV screenshot depicting the unaltered expression of *OsWRKY69* by OsWRKY53 in response to OsPep2 treatment.

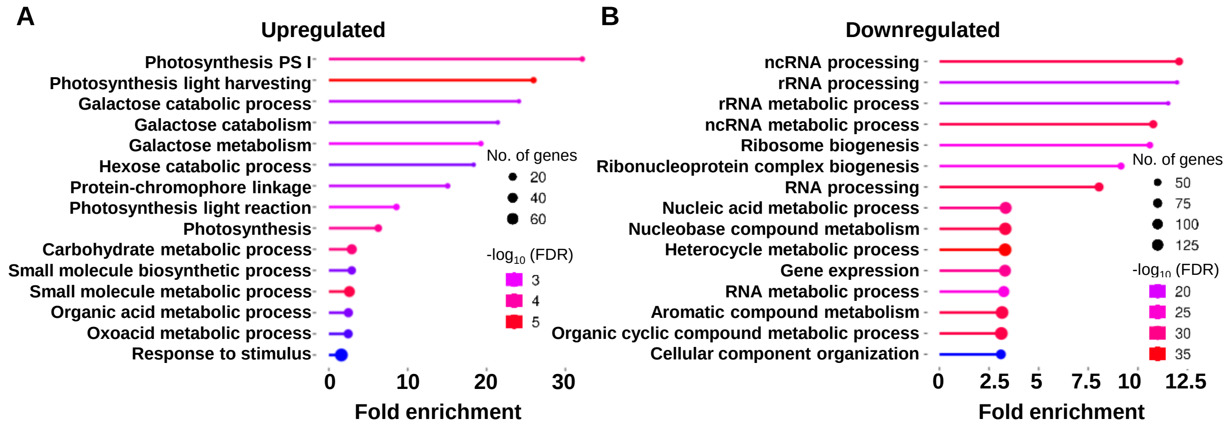


**Figure S7: Similar categories of genes are contrastingly regulated by OsPSKR and OsWRKY53**
**(A)** GO analysis of genes upregulated in OsPSKR OE as well as Oswrky53 KO. **(B)** GO analysis of genes downregulated in OsPSKR OE as well as Oswrky53 KO.
